# Heparan sulfate glycosaminoglycans mediate CXCL4 (PF4) transport across the blood-brain barrier and effects on neurogenesis

**DOI:** 10.1101/2025.04.23.650173

**Authors:** Riley R Weaver, Anna L Gray, Saudina Mateus-Gomes, Amanda JL Ridley, Sean P Giblin, Holly L Birchenough, Francis C Peterson, Kirstin O Lowe, Yui Matsubara, Iashia Z Nabi, Ingo Schiessl, Thomas A Jowitt, James E Pease, William A. Banks, Rejane Rua, Michelle A Erickson, Douglas P Dyer

## Abstract

CXCL4 (PF4) is a chemokine stored in platelets that has pleiotropic effects across biological settings. These effects include driving of inflammation and fibrosis as well as reversal of the effects of ageing. We have recently demonstrated that CXCL4 function is driven, independently of known chemokine receptors, through binding to glycosaminoglycan (GAG) side chains on proteoglycans within the cell surface glycocalyx. In this study, we have used intravital imaging and radioactive tracer studies, in combination with an exogenous inhibitor and a GAG-binding CXCL4 mutant, to demonstrate that CXCL4 can enter the brain parenchyma of mice by binding to proteoglycans within the cell surface of the endothelial glycocalyx of the blood-brain barrier (BBB). Furthermore, we have also demonstrated that CXCL4 directly promotes neurogenesis *in vitro*, which is mediated by its ability to oligomerise and bind to GAGs. These findings provide a molecular mechanism for CXCL4 uptake and function within the brain. Furthermore, these data have important implications for understanding CXCL4 during health and disease that may enable development of CXCL4-related therapeutics for inflammatory diseases and ageing.

## Introduction

CXCL4 is a chemokine that was initially identified as a platelet-derived factor that neutralizes the anticoagulant activity of heparin, and can mediate heparin’s negative side effects such as thrombocytopenia (*1, 2*). Many functions of CXCL4 have since been identified that include leukocyte chemotaxis, immunomodulation, and angiostatic activities (*3*). More recent works have shown that systemically administered CXCL4 improves cognition by mechanisms that include attenuating neuroinflammation, improving systemic immune functions, and promoting the survival or maturation of new neurons from neural precursor cells (*4–6*). Although these activities of CXCL4 have been attributed to its peripheral effects in one study (*4*), another has suggested an ability to enter and act locally within the brain (*5*). Crucially, the receptor for CXCL4 had not, until recently, been conclusively identified. Our recent work has now shown that CXCL4 interaction and cross-linking of heparan sulfate glycosaminoglycans (HS GAGs), attached to proteoglycans on the endothelial surface, are sufficient to mediate endothelial signaling and leukocyte recruitment; independently of all known chemokine receptors (*7*). Specifically, this work suggests that HS proteoglycans within the endothelial glycocalyx are in fact the receptors for CXCL4, and potentially for other chemokines too (*8, 9*). This adds to the previously understood function of chemokine:GAG interactions in facilitating their correct localisation (*10, 11*).

HS GAGs are found within the endothelial glycocalyx attached to proteoglycans, including in the brain vasculature where they are a key component of the blood-brain barrier (BBB) (*12*). HS proteoglycans also occur on the surface of a wide range of cells relevant to the brain, including stromal and immune cells (*13, 14*). Other chemokines including CCL11, CCL2, and CCL5 can cross the intact BBB (*15, 16*), and CCL2 and CCL5 do so via HS GAG-dependent mechanisms (*15*). Therefore, we posited that CXCL4 may also cross the BBB in the blood-to-brain direction through its interactions with the HS-GAGs of the brain endothelial glycocalyx and that some of the neuroprotective effects of CXCL4 could be through direct activities on brain cells.

Here, we characterized the binding interactions of wild-type recombinant mouse CXCL4 with a HS GAG surrogate, heparin octasaccharide dp8, vs. a K56E mutant mouse CXCL4 that is homologous with the human K50E mutation that has weakened GAG-binding properties (*17*). We then evaluated the blood-to-brain transport and brain retention of wild-type and K56E mutant mouse CXCL4, as well as transport from blood into other tissues using well-established, highly sensitive radiotracer assays as well as intravital microscopy. We further analysed the effects of inflammation on the BBB transport of CXCL4 using a moderate (0.3mg/kg) dose of LPS that does not induce leakage of the BBB (*18*). Finally, we evaluated the direct activities of wild-type and mutant mouse CXCL4 on neural stem cell differentiation. Overall, our results support that CXCL4 can enter the brain from the circulation by crossing the BBB through its interactions with HS GAGs of the venular endothelial glycocalyx, and also uses GAG-dependent mechanisms to stimulate differentiation of neural stem cells.

## Results

Several recent publications have demonstrated that CXCL4 enhances cognition in aged animals (*4–6*), however the underlying mechanisms producing these observations remain unknown. We have recently demonstrated that glycocalyx proteoglycans lining the brain vasculature act as the functional receptor for CXCL4 (*7*). Therefore, we hypothesised that GAG interactions may drive CXCL4 interactions with the vasculature of the brain, enabling uptake into the CNS and direct function in this location.

### Murine CXCL4 interactions with glycocalyx GAGs are driven by CXCL4 oligomerisation

We have shown that oligomerisation of human CXCL4 drives its unique high-affinity interaction with GAGs (*7, 17*). Murine and human CXCL4 are well conserved, but some differences occur in the amino acid sequence (Fig. 1A). Therefore, we assessed the biophysics and nature of the murine CXCL4:GAG interaction (Fig. 1) to enable subsequent functional studies in mice. We firstly used sedimentation velocity analytical ultra centrifugation (SV-AUC) to analyse the sedimentation co-efficient, indicative of oligomerisation status, of wild type murine CXCL4 (mCXCL4) with and without GAG (heparin dp8) (Fig. 1B and Supp. Fig. 1). This approach demonstrated that by itself mCXCL4 has a sedimentation co-efficient of 2.46s, indicating dimerization. In the presence of heparin dp8 (10:1 dp8:chemokine), mCXCL4 had a sedimentation co-efficient of 3.76s, indicating formation of a tetramer in the presence of GAG, in agreement with previous studies of hCXCL4 (*7, 19*). We then demonstrated that the K56E mutation inhibits mCXCL4 oligomerisation, just as it inhibits hCXCL4 oligomerization (*17*). Specifically, the K56E mCXCL4 mutation reduced the sedimentation co-efficient to 1.2s (indicative of a monomer) and 2.5s (indicative of a dimer) in the absence and presence of GAG (dp8), respectively.

**Figure 1.**
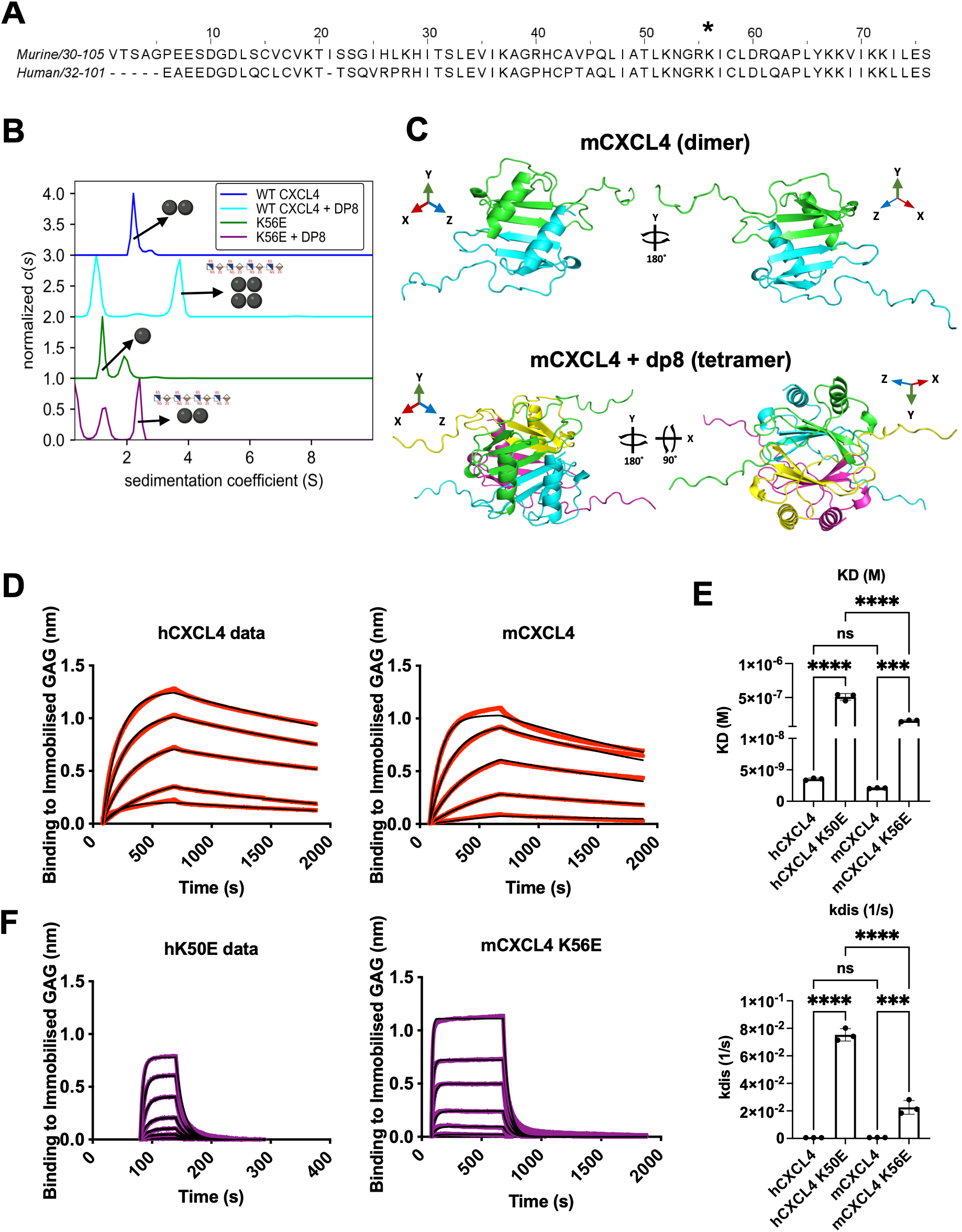
Murine CXCL4 has a high-affinity interaction with GAGs driven by oligomerization. (A) Sequence alignment on murine and human CXCL4, with human K50 highlighted to show murine K56 is the predicted equivalent residue. (B) AUC was used to determine the oligomerization status of WT and K56E mutant mCXCL4 with and without heparin DP8 GAG. (C) AlphaFold2 prediction of the structure formed by mCXCL4 in its dimeric and tetrameric forms. The top-ranking model for each was visualized in PyMol v2.5, with the dimer displayed with 180° Y-axis rotation, and the tetramer with both 180° Y-axis and 90° X-axis rotation. (D) Heparin DP8 was immobilized to a sensor to allow kinetic binding experiments with wild type human and murine CXCL4, (E) curves were then fitted to allow estimation of binding constants. (F) Kinetic binding experiments of human and mouse non-oligomerizing mutants. **p<0.01, *** p<0.001, ****p<0.0001 via Sidak’s multiple comparison test.

We then used AlphaFold to predict the structure of murine CXCL4 in the absence (dimer) and presence of GAG (tetramer) (Fig. 1C and Supp. Fig. 2). Bio-layer interferometry (BLI) interaction studies demonstrated that murine and human CXCL4 have comparable high-affinity interactions with GAGs with an overall binding affinity (K_D_) of 2.1 (± 0.1) and 3.5 (± 0.2) nM, respectively (Fig. 1D-E and Supp. Fig. 3). Furthermore, BLI confirmed that the mCXCL4 K56E mutant had a similarly reduced affinity for immobilised GAG as the human K50E equivalent and that this was driven by a much quicker off-rate of interaction as compared to the wild type version (Fig. 1E-F and Supp. Fig. 3). Crucially, we have previously shown that the hCXCL4 K50E with the same rapid off-rate cannot cross-link GAGs and therefore cannot signal through proteoglycans into endothelial cells. Taken together, these data determined that human and murine CXCL4 are very similar in the affinity and nature of their interactions with GAGs and that the K56E mouse mutation diminishes overall GAG-affinity due to reduced oligomerisation capacity and cross-linking of GAGs.

### GAG binding mediates CXCL4 transport across the mouse blood-brain barrier

Prior studies have shown that chemokines can cross the intact mouse BBB (*15, 16*), and that BBB transport involves binding of HS GAG (*15*). However, the previous studies establishing a role for CXCL4 in the CNS produced variable results on its capacity to enter and function directly in the CNS (*4–6*). Thus, we reasoned that CXCL4 may also cross the BBB via HS GAG-dependent mechanisms.

Using flow cytometry assays with fluorescently labelled CXCL4, we evaluated binding to CD45^−^CD31^+^HS^+^ brain endothelial cells in the presence and absence of exogenous heparin (20U/ml), gating detailed in Supp. Fig. 4. We found that incubation with heparin significantly reduced binding of CXCL4 to HS^+^ brain endothelial cells by approximately 90% (Fig. 2A-B). We then used highly sensitive radiotracer assays to further characterize the involvement of CXCL4 HS-GAG binding on its BBB transport. We first evaluated the labelling of WT and K56E CXCL4 with ^125^I, and found that specific activity of the labelled proteins was equivalent, with both ^125^I-labeled proteins migrating on SDS-PAGE at molecular weights that were the same as the unlabeled proteins (Supp. Fig. 5). No radiolysis products or aggregates were detected for either protein form. Our initial studies of *in situ* brain perfusions (Fig. 2C) in adult CD-1 mice showed that wild-type mouse CXCL4 crossed the intact BBB in situ at a rate of 2.894±0.8363 ul/g/min (R^2^=0.5994, p=0.0086 vs. zero slope) that was significantly reduced to 0.1415±0.8363 ul/g/min (R^2^=0.1548, p=.2606 vs. zero slope) by co-perfusion with 20U/ml heparin (p=0.0058, F=10.10 (1,16)) (Fig. 2D). These data confirm that CXCL4 can bind to HS^+^ endothelial cells and rapidly cross the BBB in a GAG-dependent manner.

**Figure 2.**
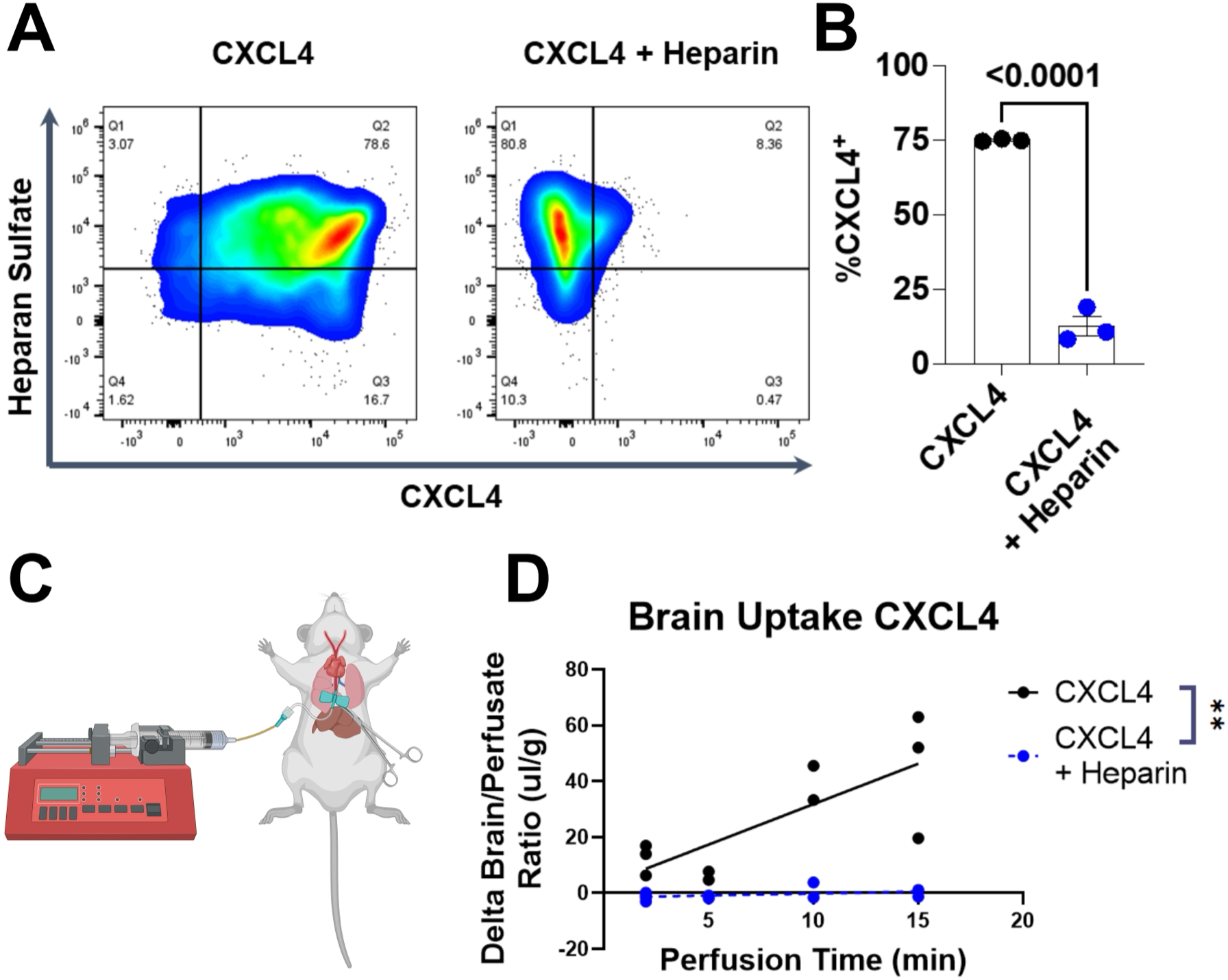
HS-GAGs mediate interactions of CXCL4 with brain endothelial cells and transport across the BBB *in situ*. A) Mouse brain endothelial cells (CD45-CD31+) were analysed using flow cytometry for their ability to bind to fluorescently labelled mCXCL4 via HS in the presence and absence of 20U/ml heparin. B) Quantification of the percentage of CXCL4+ cells in the presence and absence of 20U/ml heparin. C) Illustration of the *in situ* brain perfusion setup. D) *In situ* BBB transport of ^125^I-CXCL4 in the presence or absence of 20U/ml heparin, **p< 0.001 indicating significant difference of slopes. Figure generated with BioRender.

We next determined whether there were differences in the BBB transport of wild-type and the HS-binding mutant K56E mouse CXCL4 *in vivo*, using the multiple-time regression analysis technique to calculate *in vivo* chemokine transport rates across the BBB (Fig. 3A). The blood clearance of WT and K56E mCXCL4 significantly differed (p< 0.0001, F=12.23 (3,30)) (Fig 3B), and half-life of clearance for WT was 8.606 min vs. 2.576 min for K56E. The transport rate of WT mCXCL4 across the mouse BBB (Fig 3C) was 1.114 ± 0.1236 ul/g-min (R^2^= 0.8354) vs. 2.736 ± 0.4754 (R^2^=0.6744); the slopes of WT vs. K56E significantly differed (p=0.0006, F=14.64 (1,32)). We did not observe significant or apparent differences in BBB transport rates of WT or K56E mCXCL4 in male vs. female mice. Additionally, we calculated transport rates of WT and K56E mCXCL4 in peripheral tissues from the same mice. In peripheral organs, the vascular space-corrected transport rates of CXCL4 were all significantly non-zero, and ranged from 1.683 ul/g/min in cecum to 127.2 ul/g/min in the liver (Table 1). Surprisingly, the transport rates and vascular binding (Vi) of the mCXCL4 K56E mutant were higher in the brain and in peripheral tissues, except for spleen.

**Figure 3.**
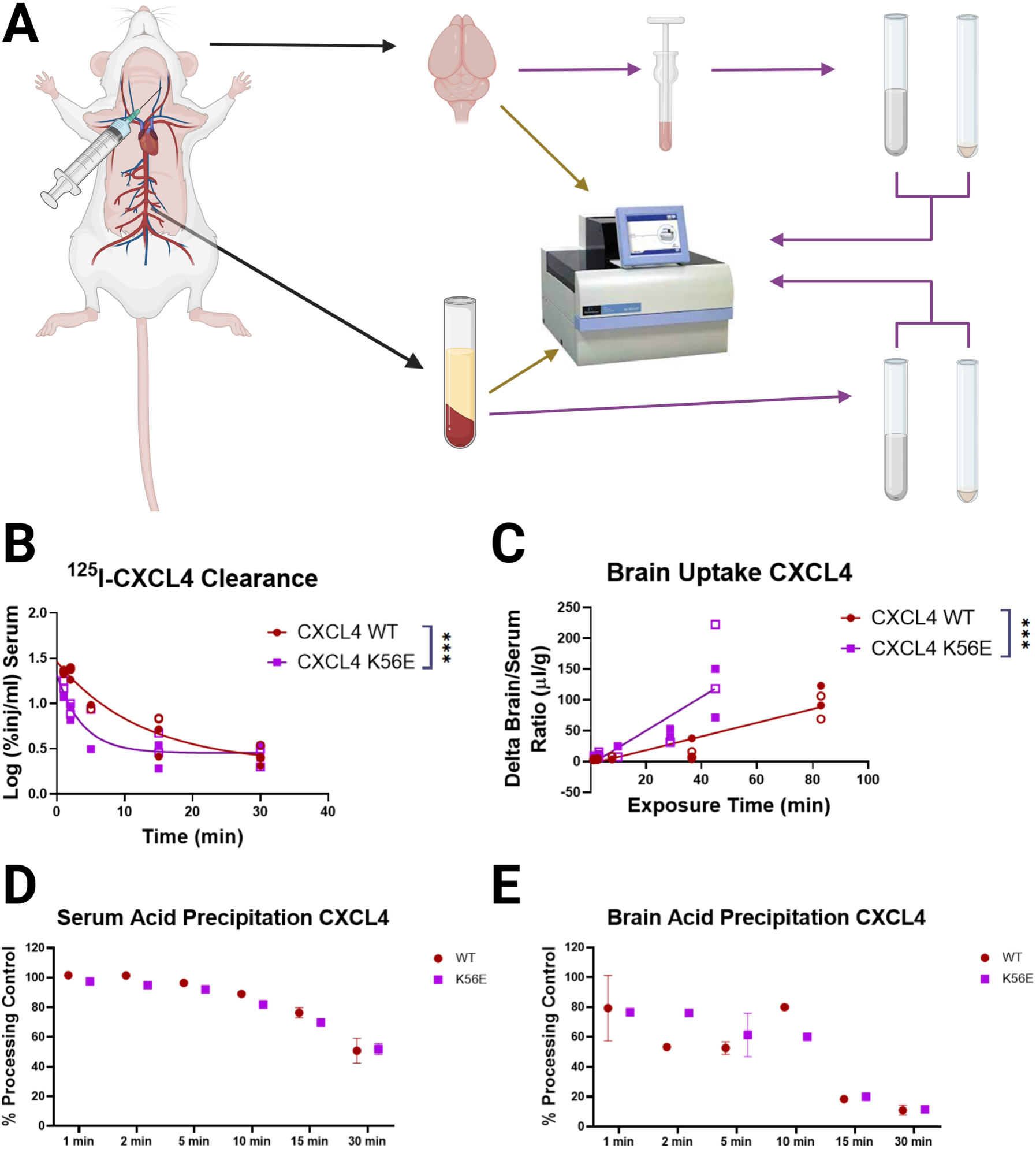
Transport of wild-type and K56E mutant ^125^I-CXCL4 across the BBB in vivo, and stability in blood and brain. A) Illustration of *in vivo* transport study design. Black arrows depict the collection of mouse brain and serum, which are then either counted (brown arrows), or processed for acid precipitations prior to gamma counting (purple arrows). B) Clearance of ^125^I-CXCL4 WT and K56E mutant forms from blood over a 30 min circulation time following IV injection in vivo. ***p<0.001 in comparison of fits test, indicating that the curves have significantly different fits. C) Multiple-time regression analysis of ^125^I-CCL4 WT and K56E mutant brain uptake ***p< 0.001 for the comparison of slopes of each line. In B and C, open shapes are female mice, and closed shapes are male mice. D) Stability of ^125^I-CXCL4 WT and K56E forms over time in serum and E) in brain, normalized to a processing control for each tissue. Figure generated with BioRender.

**Table 1.**
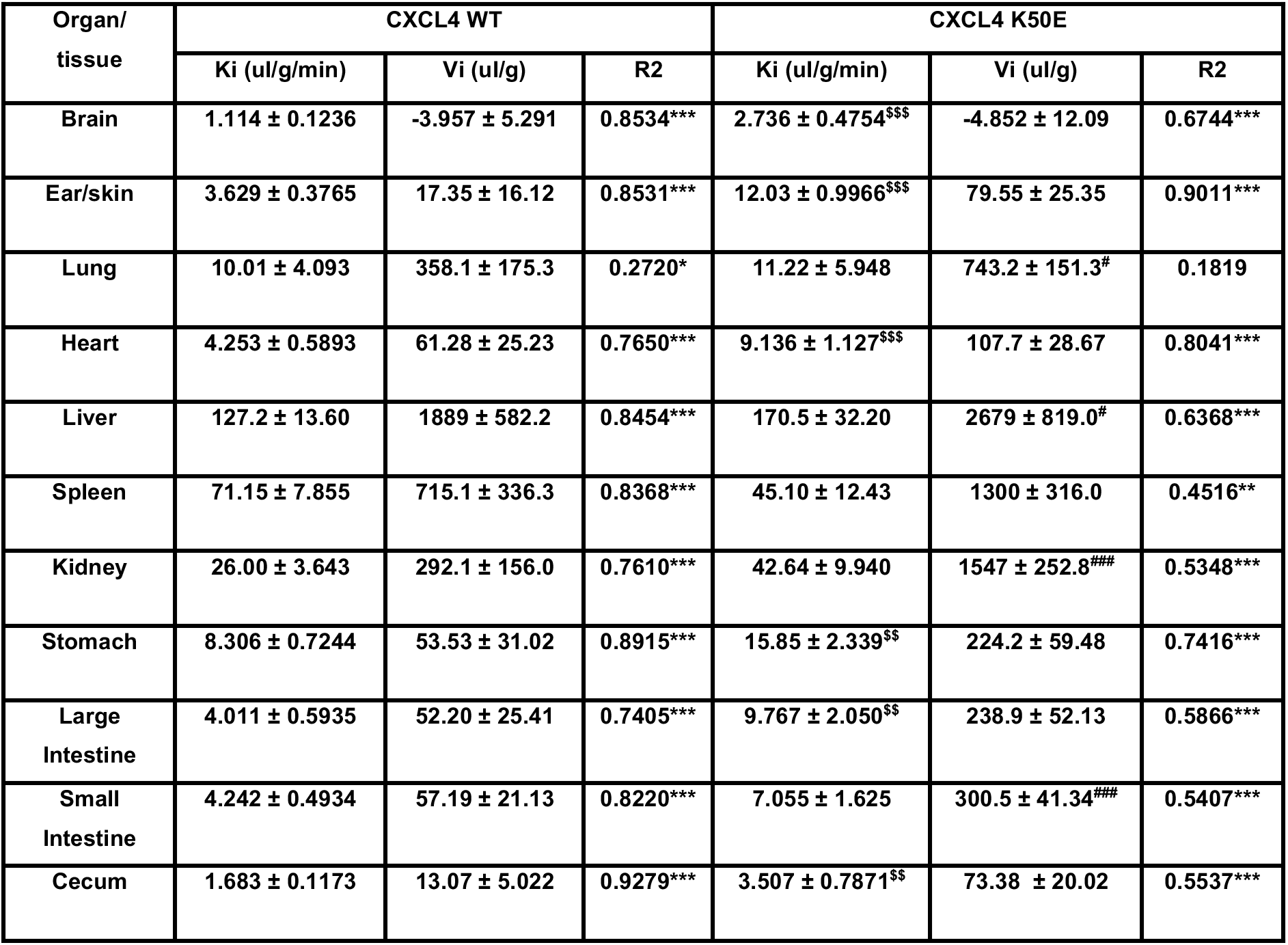
Transport rates (Ki) and vascular binding (Vi) of ^125^I-CXCL4 WT and K56E mutants in different tissues, corrected for 99mTc-albumin vascular space. *p < 0.05, **p<0.01, ***p<0.001, significantly non-zero slope. #p<0.05, ###p<0.001, difference of intercept vs. WT. $$p<0.01, $$$p<0.001, difference of slope vs. WT. Intercept differences aren’t compared when slopes significantly differ.

| Organ/<br>tissue | CXCL4 WT |  |  | CXCL4 K50E |  |  |
| --- | --- | --- | --- | --- | --- | --- |
|  | Ki (ul/g/min) | Vi (ul/g) | R2 | Ki (ul/g/min) | Vi (ul/g) | R2 |
| Brain | 1.114 ± 0.1236 | -3.957 ± 5.291 | 0.8534*** | 2.736 ± 0.4754<sup>\$\$\$</sup> | -4.852 ± 12.09 | 0.6744*** |
| Ear/skin | 3.629 ± 0.3765 | 17.35 ± 16.12 | 0.8531*** | 12.03 ± 0.9966<sup>\$\$\$</sup> | 79.55 ± 25.35 | 0.9011*** |
| Lung | 10.01 ± 4.093 | 358.1 ± 175.3 | 0.2720* | 11.22 ± 5.948 | 743.2 ± 151.3<sup>#</sup> | 0.1819 |
| Heart | 4.253 ± 0.5893 | 61.28 ± 25.23 | 0.7650*** | 9.136 ± 1.127<sup>\$\$\$</sup> | 107.7 ± 28.67 | 0.8041*** |
| Liver | 127.2 ± 13.60 | 1889 ± 582.2 | 0.8454*** | 170.5 ± 32.20 | 2679 ± 819.0<sup>#</sup> | 0.6368*** |
| Spleen | 71.15 ± 7.855 | 715.1 ± 336.3 | 0.8368*** | 45.10 ± 12.43 | 1300 ± 316.0 | 0.4516** |
| Kidney | 26.00 ± 3.643 | 292.1 ± 156.0 | 0.7610*** | 42.64 ± 9.940 | 1547 ± 252.8<sup>###</sup> | 0.5348*** |
| Stomach | 8.306 ± 0.7244 | 53.53 ± 31.02 | 0.8915*** | 15.85 ± 2.339<sup>\$\$</sup> | 224.2 ± 59.48 | 0.7416*** |
| Large Intestine | 4.011 ± 0.5935 | 52.20 ± 25.41 | 0.7405*** | 9.767 ± 2.050<sup>\$\$</sup> | 238.9 ± 52.13 | 0.5866*** |
| Small Intestine | 4.242 ± 0.4934 | 57.19 ± 21.13 | 0.8220*** | 7.055 ± 1.625 | 300.5 ± 41.34<sup>###</sup> | 0.5407*** |
| Cecum | 1.683 ± 0.1173 | 13.07 ± 5.022 | 0.9279*** | 3.507 ± 0.7871<sup>\$\$</sup> | 73.38 ± 20.02 | 0.5537*** |

Additional studies characterized the *in vivo* stability of CXCL4 by acid precipitation. As degradation products of iodinated proteins are rapidly cleared and generally cross the BBB poorly (*20*), we have found that multiple-time regression analysis often adequately corrects for degradation, with rates of uptake reflecting the transport of intact protein. The purpose of acid precipitation experiments was to compare degradation over time of the WT vs. K56E mutant CXCL4 in brain and blood. Figure 3D and E show that the stability of WT and K56E CXCL4 over time is approximately equivalent in blood and brain, respectively. Further, the degradation of CXCL4 in brain was higher than that of blood at equivalent time points, indicating that CXCL4 is rapidly degraded in brain post-uptake.

### CXCL4 oligomerisation regulates chemokine retention within the cerebral vasculature

We next hypothesised that the faster rates of uptake of the mCXCL4 K56E mutant into the brain, compared to wild type, are due to the more rapid dissociation kinetics of CXCL4 from bound GAGs (Fig. 1). Thus, the mutant can still bind to the surface proteoglycans but is released from the vasculature into the parenchyma more rapidly.

To test this hypothesis, we used radiotracer assays to quantify the percentage and total amount of tracer that is bound/internalized by the brain vasculature vs. that which traverses the brain vasculature into brain parenchyma. Figure 4A and B show that there are time-dependent differences in the relative vascular and parenchyma partitioning of CXCL4 WT and K56E mutant. At 5 minutes post-injection, there were no apparent differences in the vascular-parenchyma partitioning of WT vs mutant CXCL4. However, at 15 minutes post-injection, there was significantly less K56E mutant in the vascular fraction, and more in the parenchymal fraction. Also, by 15 minutes, most of the CXCL4 of both WT and K56E had fully crossed the BBB and entered brain parenchyma. Figures 4C and D show the tissue/serum ratios of brain vascular and parenchymal uptake of CXCL4 WT and K56E proteins, calculated as percentages of whole brain uptake levels that were quantified prior to separating the vascular and parenchymal fractions and corrected for residual albumin vascular space. Results in Figures 4 C-D show that there is a higher amount of uptake overall for K56E in both capillary and parenchymal fractions by 15 minutes, reflecting its faster rate of transport. These data support that the faster K56E transport rate could be due to the ability to traverse the endothelial cells into brain parenchyma more rapidly than the WT CXCL4, as has been shown in the lung for CXCL8 (*21*).

**Figure 4.**
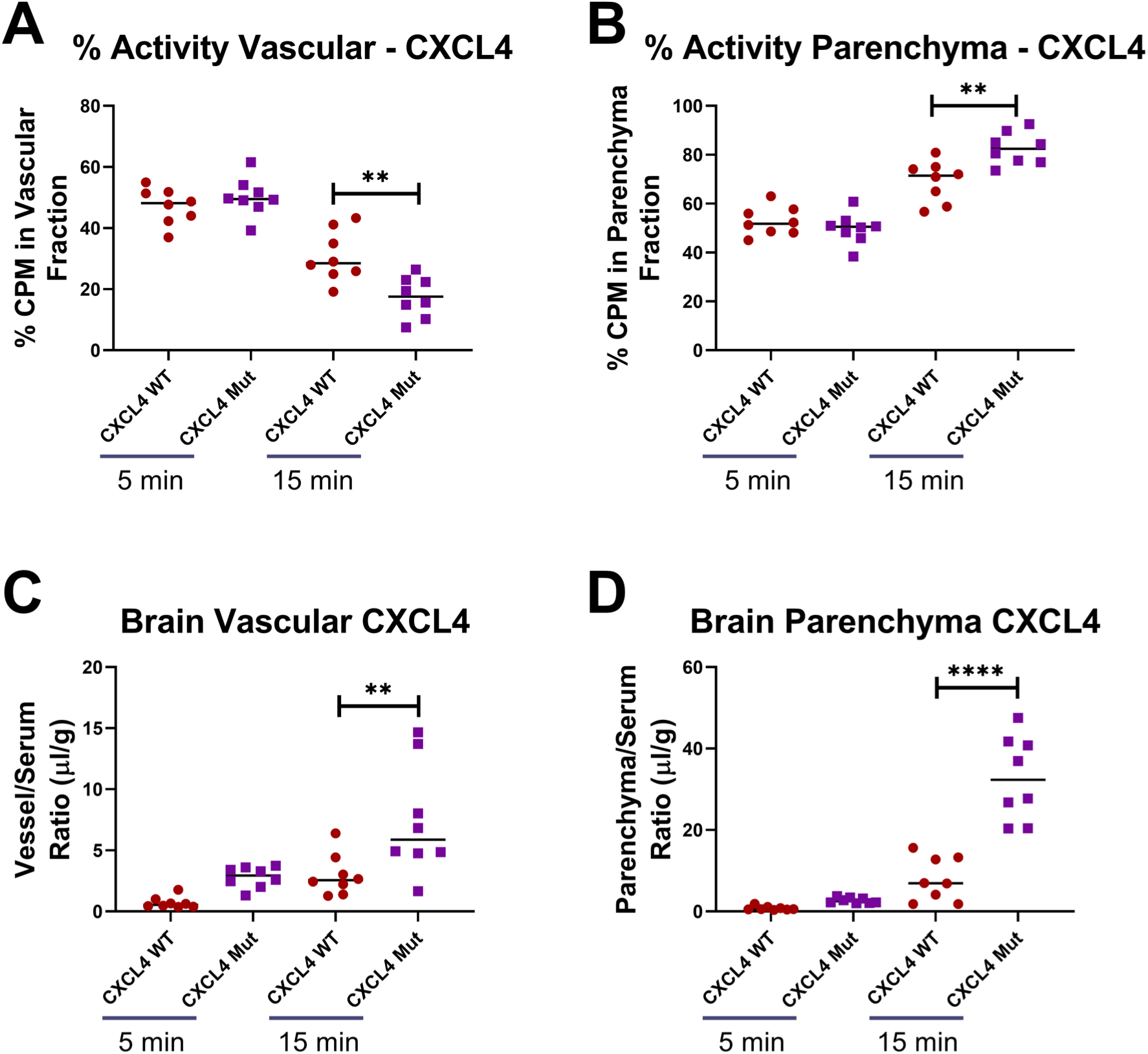
CXCL4:GAG binding kinetics regulate movement through the vascular endothelial cells into the brain. The percentage of ^125^I-CXCL4 WT or K56E CPM that partitions into the vascular A) vs. parenchymal B) fractions after 5 or 15 minutes of circulation time is shown, **p<0.01 via Sidak’s multiple comparison test. The fraction percentages and CPM values of whole brains pre-processing were used to calculate vessel/serum ratios C) or parenchyma/serum ratios D); **p<0.01, ****p<0.0001.

### CXCL4 brain uptake occurs predominantly across veins, venules, and capillaries and is rapidly cleared/degraded after it enters the brain

To further evaluate the transport of CXCL4 across the mouse BBB, we used intravital microscopy (Fig. 5A) to determine the localization of intravenously injected fluorescently labelled WT CXCL4 in the brain vasculature and brain parenchyma post-uptake. Fig 5B-D shows that fluorescently labelled CXCL4 selectively attached to large veins of the pial surface, but not arteries, 60 mins after being injected intravenously. Fig. 5D further shows the binding/co-localization of CXCL4 with wheat-germ agglutinin (WGA) positive capillaries 60 mins after being injected intravenously. Figure 5E shows the apparent uptake of WT CXCL4 into neuronal cells 150 minutes after injection.

**Figure 5.**
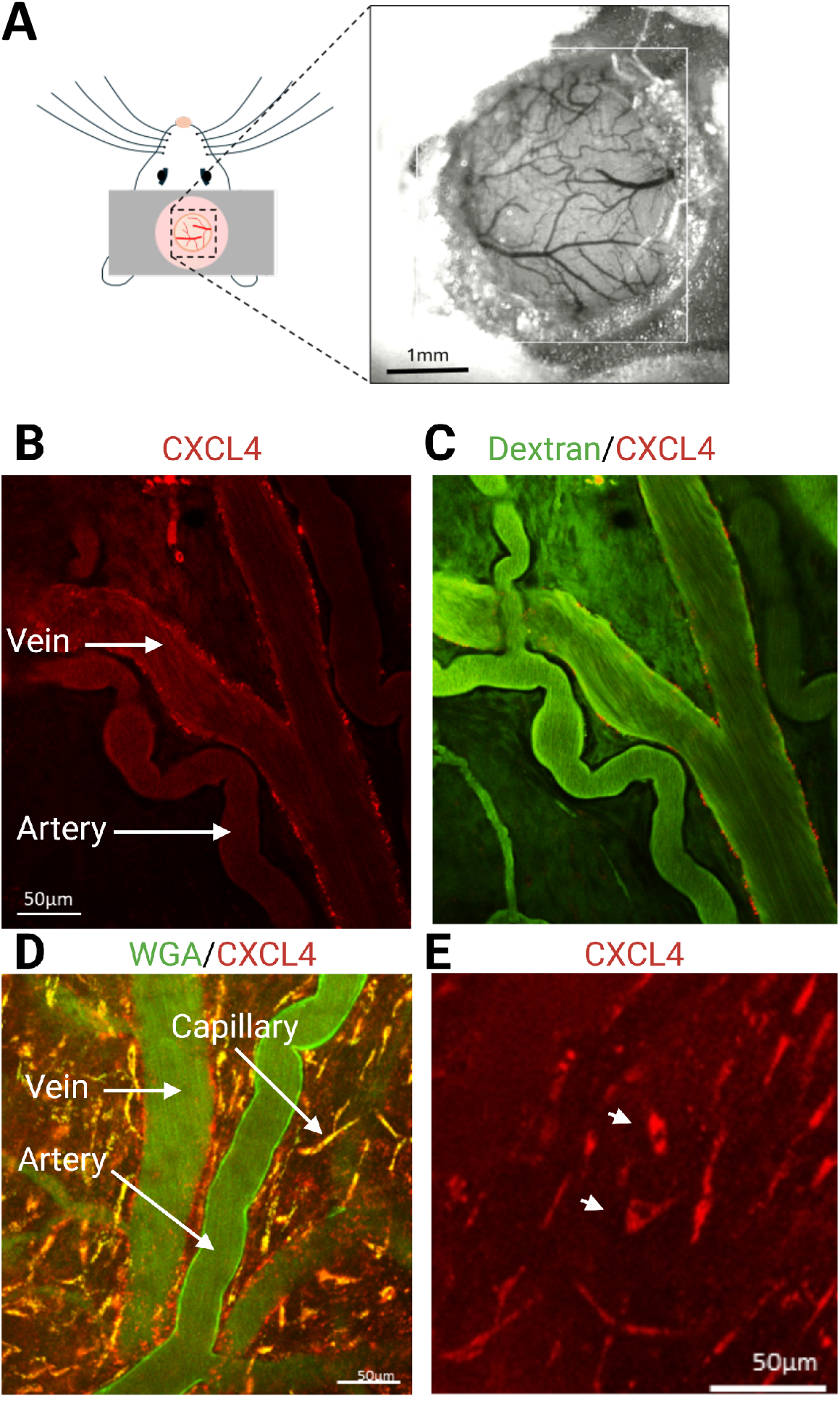
Intravital imaging of IV-injected WT fluorescently labeled CXCL4 (red) reveals uptake into the brain. A) Schematic of the imaging from the pial surface. CXCL4 fluorescence alone B) or merged with dextran fluorescence (green) to mark the vasculature C) after an 60 min circulation time shows that CXCL4 is predominantly bound to veins but not arteries. D) CXCL4 (red) and WGA (green) dual imaging after 60 min circulation time shows another example of CXCL4 localized in veins and capillaries, but not arteries. E) CXCL4 after 150 min circulation time enters brain cells with neuronal morphology (white arrow heads).

We also used fluorescent and radiochemical assays to evaluate the clearance of CXCL4 from the brain. Using an *in vivo* radiotracer assay, we found that ICV injected CXCL4 WT or K56E mutant did not show statistically significant or apparent differences in the rate of brain clearance; the half-life of the pooled slopes was 27.24 minutes (Fig 6A). Intravital microscopy assays corroborated findings from radiochemical assays, showing that brain uptake of intravenously injected WT CXCL4 peaked around 60 min and was mostly cleared from the brain parenchyma by 180 min (Fig 6B), although some CXCL4 was retained and remained associated with venules (Fig 6C-D).

**Figure 6.**
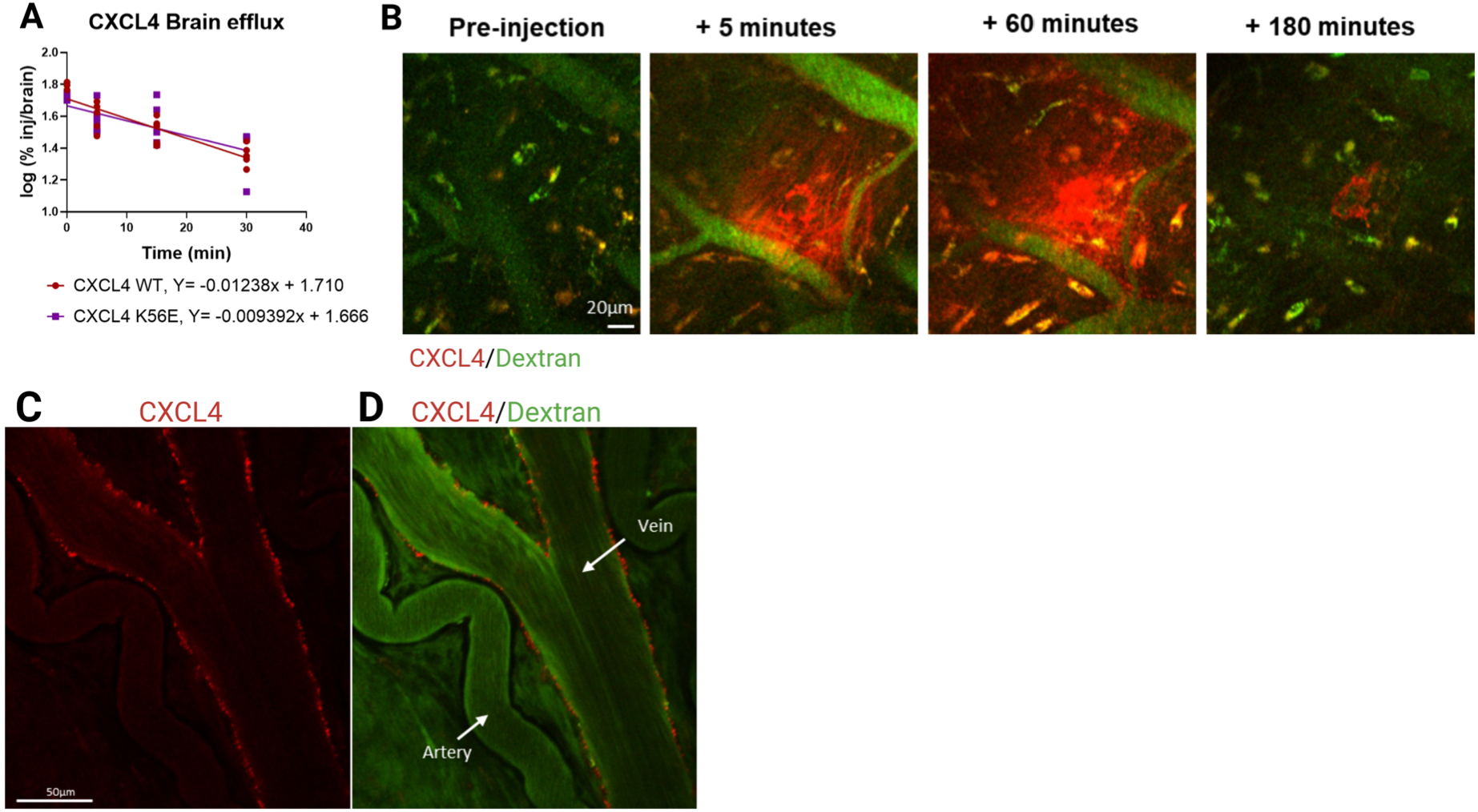
CXCL4 is rapidly cleared from the brain. A) ^125^I-CXCL4 WT or mutant forms were injected ICV into CD-1 mice and brain clearance was measured. No difference in clearance was observed for the two CXCL4 forms. B) Brain clearance of IV-injected, fluorescently labeled WT CXCL4 was visualized with intravital microscopy up to 180 minutes post-injection. C and D) Intravital brain images of IV-injected WT CXCL4 (red) and dextran (green) 150 minutes post-injection showing that CXCL4 remains bound to the veins but not the arteries.

### Moderate, acute systemic inflammation increases CXCL4 transport across the BBB

We next investigated whether inflammatory status could alter the uptake of CXCL4 into the brain. To evaluate the effects of systemic inflammation on CXCL4 transport, we treated CD-1 mice with a single dosage of 0.3mg/kg LPS, and evaluated the blood-to-tissue transport of WT CXCL4 *in vivo* and *in situ*. This dose of LPS causes weight loss and sickness behaviours but does not induce significant BBB leakage at 24 hours post-injection (*18*). The LPS treatment paradigm was secondarily chosen to compare to prior studies of CCL2 and CCL5, which showed multi-fold increases in BBB transport *in situ* following LPS treatment (*15*). *In vivo* multiple-time regression analysis studies showed that LPS significantly accelerated the clearance of CXCL4 from blood (p= 0.0285, F=3.851 (3,17)), with the half-life of clearance of vehicle being 6.996 min, and the half-life of clearance for LPS treatment 2.196 min (Figure 7A). Figure 7B shows that the albumin vascular space-corrected transport rates of CXCL4 in saline vs. LPS groups are similar, however the Y-intercept reflecting the initial vascular binding is significantly higher in LPS-treated mice (p= 0.0243, F=5.937 (1,20)). We similarly found that CXCL4 vascular binding in most peripheral organs was elevated with LPS, whereas the rates of uptake were unaffected (Figure 7C-G). We also observed that LPS increased the brain uptake of CXCL4 following a 10 min in situ brain perfusion (Fig. 7H), however the magnitude of increase was much less than that observed for CCL2 and CCL5, indicating that the inflammatory regulation of chemokine transport is selective rather than generalized.

**Figure 7.**
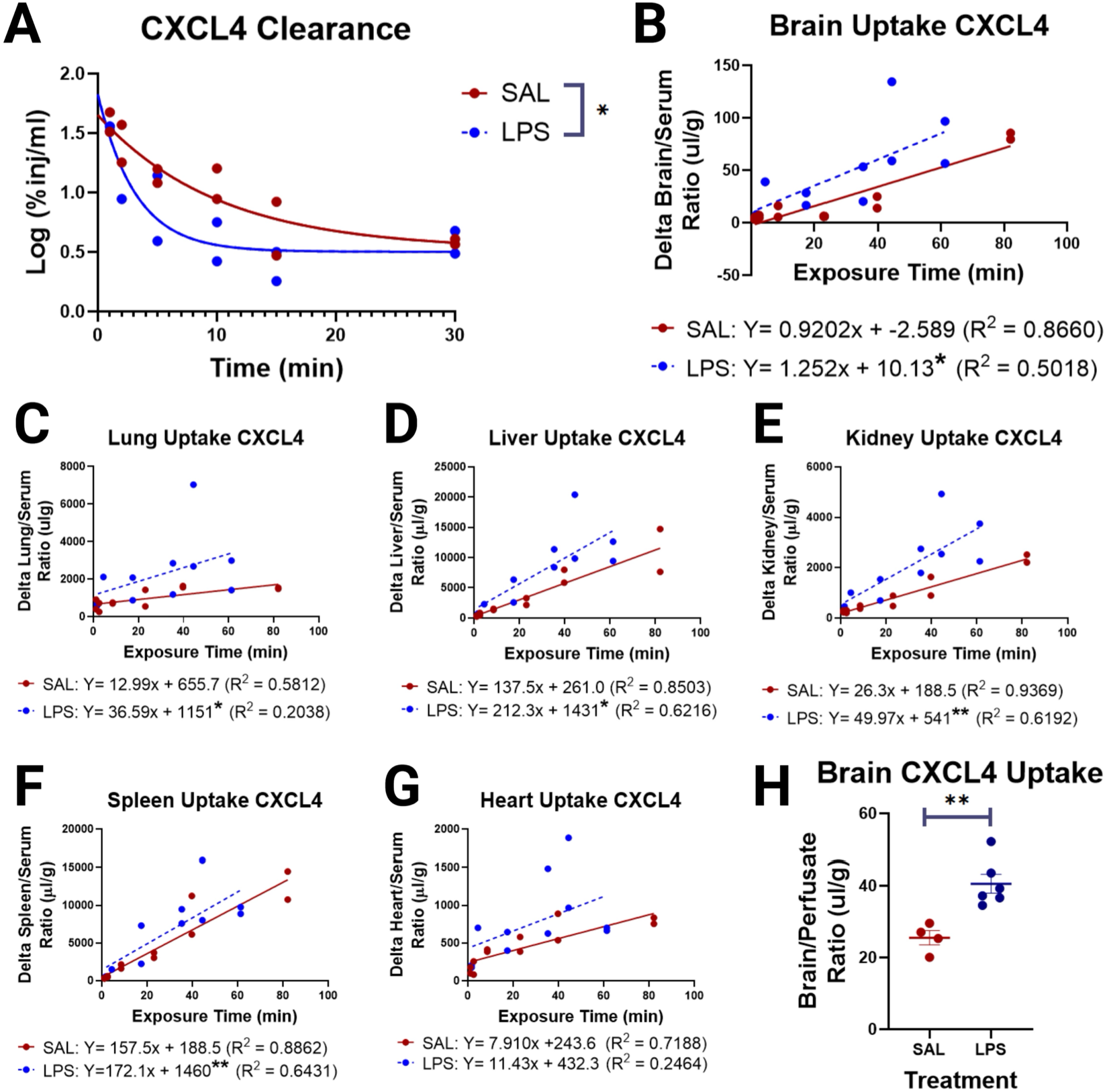
LPS-induced systemic inflammation enhances WT ^125^I-CCL4 serum clearance. A), BBB transport B), and uptake in to lung C), liver D), Kidney E), Spleen F), and Heart G) in vivo. In situ brain perfusions are shown in H). *p<0.05, **p<0.01. Statistics indicated in B-G are comparing the vascular binding (Y intercept) between saline (SAL) and LPS administered groups.

### BBB transport and vascular/parenchymal partitioning is not significantly altered in aged mice

Circulating CXCL4 was shown to protect and improve neurologic functions, particularly in older mice (*4–6*). However, recent work has also identified that age-associated changes in glycocalyx composition can impact BBB functions (*12*), although the impact on HS-dependent BBB transport is unknown. Therefore, we next determined whether brain uptake or vascular/parenchymal partitioning of CXCL4 is altered with normative aging in C57BL/6J mice. Supplementary Figure 6A shows that there were no apparent differences in brain uptake of WT ^125^I-CXCL4 after 15 minutes of circulation time in young (2 month) vs. old (22 month) mice. Supplementary Figure 6B shows that there is no significant whole brain BBB leakage to ^99m^Tc-albumin in young vs. old mice. Supplementary Figures 6C-F show that the partitioning of ^125^I-CXCL4 into brain vascular and parenchymal fractions was unaltered with aging. Therefore, CXCL4 transport across the BBB into brain parenchyma is unaltered with aging.

### CXCL4 regulates neural stem cell differentiation in vitro

Prior work has shown that systemic CXCL4 impacts neurogenesis by increasing the numbers of neurons in the hippocampus of young and old mice *in vivo*, which is associated with improved cognitive function (*6*). Having shown that CXCL4 is transported into the brain parenchyma in a GAG-dependent fashion, we next wished to determine whether CXCL4 can directly exert its effects on neurogenesis, and whether HS-GAG binding is involved. As expected, after a six-day incubation of mouse neurospheres with WT CXCL4, we found an increased density of adherent, differentiated cells and a concomitant reduction of the density of undifferentiated neurospheres (Fig. 8). In contrast, K56E CXCL4, which does not oligomerise and has reduced GAG-binding capacity, had no effect on neurosphere differentiation. These data confirm that CXCL4 could contribute to neurogenesis through direct interactions with neural stem cell populations, and that GAG-binding is crucial to CXCL4-mediated neurogenesis.

**Figure 8.**
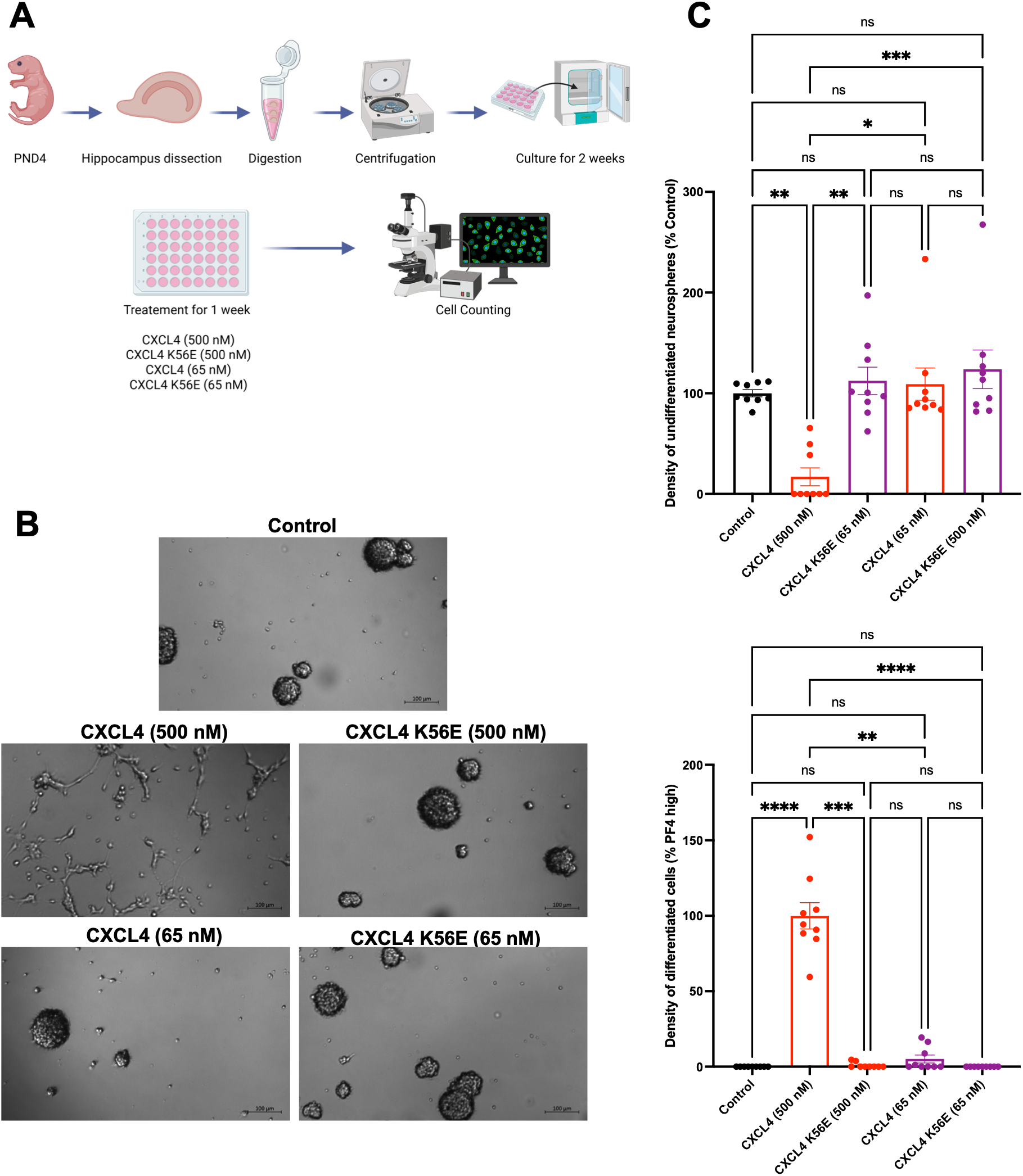
CXCL4 oligomerisation mediates its ability to promote neurogenesis. A) Primary neurospheres were cultured for 6 days with PF4 or PF4 mutant (at high and low doses), and undifferentiated floating versus differentiated cells were imaged (B) and quantified (C) on day 6. Bar graphs show the mean ± SEM, and asterisks denote statistical significance (ns=non-significant, *p < 0.05; **p < 0.01; ***p < 0.001; ****p < 0.0001 Kruskal-Wallis test followed by Dunn’s multiple comparison test). Data are representative of 3 independent experiments with 9 samples per group.

## Discussion

CXCL4 (PF4) has long been an enigma in the chemokine field, possessing the structural characteristics of a chemokine but not directly inducing immune cell chemotaxis in simplified systems. However, *in vivo* studies of CXCL4 knock out mice show that CXCL4 regulates the recruitment of immune cells in multiple inflammatory contexts, such as neutrophil trafficking to lungs after acute injury and T cell trafficking to the CNS during malarial infection (*22, 23*). Furthermore, a consistent finding of CXCL4 function from these approaches is that it regulates fibrosis in many tissues and has been proposed as a link between inflammation and fibrosis (*24*). In addition to this role during inflammation and immune cell recruitment, a series of recent papers have proposed that CXCL4 can have beneficial effects, such as enhancing neurogenesis and cognition in aged mice (*4–6*).

Despite the importance of CXCL4 function across biological contexts, the molecular mechanism (i.e. receptor) that regulates CXCL4 functions has remained unknown (*25*). Whilst some historical studies have suggested a very weak signalling capacity for CXCL4 through CXCR3b and CCR1, more recent studies have not been able to reproduce some aspects of these findings (*26–28*). Our lab recently proposed that the receptor for CXCL4 is in fact the HS proteoglycans such as those found on endothelial cells, and increasingly other cells too, via binding and remodelling of their GAG sugar side chains (*7*). The experiments detailed in this manuscript further support that it is the GAG binding capacity of CXCL4 that is critical to its function. This builds on a series of consistent findings across tissues and inflammatory contexts that CXCL4 function is GAG-binding dependent (*25, 29, 30*). Specifically, use of exogenous heparin as an inhibitor of binding to endogenous HS-GAGs, in addition to a mutant of CXCL4 (K56E) that does not cross-link GAGs, has demonstrated that GAG-binding is critical to the uptake of CXCL4 into the brain and its subsequent effects on neurogenesis (Fig. 9). Our findings thus provide a molecular mechanism for how circulating CXCL4 could enter the brain and directly produce beneficial CNS effects.

**Figure 9.**
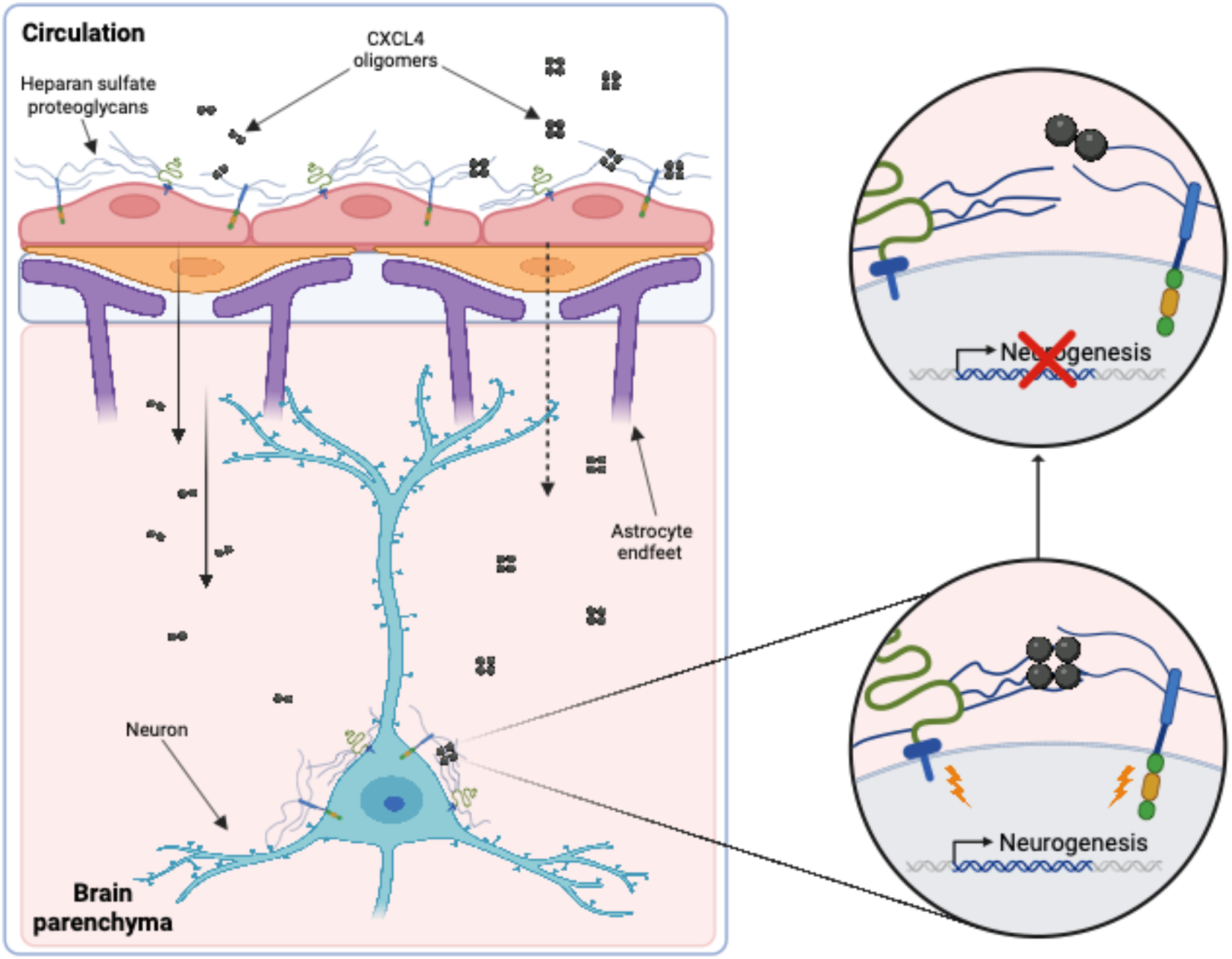
Cell surface proteoglycans mediate CXCL4 transport into the CNS and effects on neurogenesis. CXCL4 present in the circulation can bind to HS-GAGs on brain endothelial cells, be transported across the blood-brain barrier, and promote neuronal differentiation in a proteoglycan-dependent fashion.

Although prior studies in addition to this one support that HS-GAGs have BBB transporter functions (*15, 31*), the precise mechanisms by which they function as BBB transporters require further characterization. HS-GAGs likely act in tandem with their membrane-bound core proteins (i.e. syndecans and glypicans) to facilitate internalization of bound ligands and transcytosis across the BBB. Whereas HS proteoglycan (HSPG)-mediated endocytosis has been characterized in cell lines (*32, 33*), it is not yet known which HSPGs predominantly mediate BBB transport of chemokines. RNA sequencing methodologies have, however, identified unique profiles of syndecans and glypicans that are expressed in mouse and human brain microvessels: syndecans 2-4 and glypicans 1 and 3-6 at lower abundances have been detected (*34*). Interestingly, syndecan-1, which is a conventional marker of endothelial glycocalyx shedding (*35*), is not detected or detected at very low levels in most RNA sequencing studies of brain microvessels (*34, 36, 37*). Additionally, the composition of HS-GAG chains regulates ligand binding (*38*), and so alterations in HS-GAG composition at the BBB that are tissue and/or disease-specific could subsequently alter HSPG ligand binding and transport functions.

One surprising result was that the GAG-binding K56E mutant of CXCL4 demonstrated an increased rate of transport across the BBB. Although this was unexpected, a prior study *in vivo* has shown that chemokine mutations that reduce GAG binding can result in increased biological activity (*21*). Our studies evaluating uptake/partitioning of CXCL4 in the brain parenchyma and vasculature showed that the K56E mutation does not significantly affect vascular vs. parenchymal partitioning 5 minutes after injection, but by 15 minutes more K56E mutant CXCL4 had entered the parenchyma. This suggests that the faster rate of brain uptake of the K56E mutant may be due, in part, to faster transit from the endothelial cell into brain parenchyma. A faster transit rate could be due to reduced oligomerization capacity of K56E, which could result in different transporter confirmations. Alternatively, the higher dissociation constant of K56E could also contribute to a higher efficiency ligand release from the HSPG transporter.

Overall, a wider understanding of the presence and potential signalling capacity of HS proteoglycans for CXCL4 across cells of the CNS is needed. Future studies will need to address whether other chemokines that can cross-link GAGs also have the capacity to signal through proteoglycans on the cell surface. For example, we have previously demonstrated that both CCL2 and CCL5 are transported across the BBB by heparan sulfate proteoglycans (*15*). However, CCL5, but not CCL2, is able to cross-link GAGs (*8*). Therefore, it may be that CCL2 and CCL5 play different GAG-dependent roles once they have entered the brain.

In addition to their established function of transporting components from the circulation into the brain, endothelial proteoglycans are also a key component of the glycocalyx (*39*). The glycocalyx is a pericellular layer surrounding all cells to varying degrees, but is particularly prevalent on the luminal surface of vascular endothelial cells (*33*). In the context of the brain, a number of studies have shown that proteoglycans and the wider glycocalyx are critical to forming and maintaining functions of the BBB (*12, 40, 41*). Our data demonstrated an overlap with the glycocalyx binding lectin, WGA (*42*), and CXCL4 binding within the capillaries of the brain. This not only localises where CXCL4 transport may occur but also suggests that CXCL4 may regulate glycocalyx and BBB function through its ability to bind and remodel glycocalyx components as well as signalling into endothelial cells.

Our data also begin to address how CXCL4 mediates its beneficial effects on neurogenesis via oligomerisation capacity and subsequent GAG binding. Importantly there is an extensive literature that has detailed the presence of HS GAGs on the surface of neurons and our intravital imaging data suggested binding of CXCL4 to neuronal structures (Fig. 4E). Furthermore, genetic polymorphisms in genes that regulate proteoglycan synthesis are associated with a range of developmental and inflammatory diseases (*43*). This suggests these cells have the capacity to transduce CXCL4 signals through their cell surface proteoglycans. It will be key in future studies to expand on this to fully determine the molecular function, and interrogate the apparent contradiction, of CXCL4 being a pro-inflammatory factor and potentially having anti-ageing effects by promoting cognition.

In conclusion our data provides a molecular mechanism for previously detailed observations of CXCL4 entry, and function within, the CNS. This study also provides further evidence of the function, beyond physical barrier formation, of proteoglycans within the cell surface glycocalyx.

## Materials and Methods

### Materials and mice

For experiments performed in Manchester, up to four C57BL/6J mice were housed in cages of up to four in a 12 hr light/dark cycle, with free access to food and water. All experiments were carried out following ethical approval from The University of Manchester under licence from the UK Home Office (Scientific Procedures Act 1986). For experiments performed in Seattle, male and female CD-1 mice were purchased from Charles River Laboratories (Seattle, WA) and male and female C57BL/6J mice were purchased from The Jackson Laboratory (Sacramento, CA). The mice were housed on a 12/12-hour light/dark cycle, and received *ad libitum* food and water. Experiments were conducted when the mice reached 10-12 weeks of age, or 22 months of age. All studies were approved by the Institutional Animal Care and Use Committee of the Veterans Administration Medical Center and performed in a facility accredited by the Association for Assessment and Accreditation of Laboratory Animal Care. Blinding and randomisation was used where possible and POWER calculation analysis was used to design experiments. All chemokines were purchased from Protein Foundry and dp8 and heparin GAGs were purchased from Iduron.

### Analytical ultra-centrifugation analysis of CXCL4 oligomerisation

Wild type human CXCL4, human CXCL4 K50E, mouse wild type CXCL4 or mouse CXCL4 K56E were re-suspended in PBS to a final concentration of 7 µM either alone or in the presence of heparin dp8 at a ratio of 1:10 (chemokine:GAG). Samples were loaded into 2-sector cells with PBS as a reference and centrifuged at 50,000 rpm in a 4-hole An60Ti rotor monitoring the absorbance at 230 nm until sedimentation was reached. The time-resolved sedimenting boundaries were analysed using Sedfit (*44*). The resulting profiles are shown in Gussi (*45*).

### In silico structure modelling of murine CXCL4 dimers and tetramers

The dimeric and tetrameric structures of murine CXCL4 were modelled by AlphaFold2 (DeepMind, EMBL-EBI) using the ColabFold v1.3 interface as described (*46, 47*). Briefly, target amino acid sequences (Uniprot ID: Q9Z126 residues 30-105) were uploaded as PDB70 templates and multiple sequence alignments (MSA) were performed by MMSeqs2 against Uniref100 and environmental structure libraries with unpaired and paired sequences. Structure modelling for homo-oligomers was performed with model type AlphaFold2_multimer_v3 for complex prediction as described (*48*), with 12-iterances of model recycling, a stop-tolerance of t=0.0 for multimers with 200 maximum iterations and pairing strategy set to “greedy”. Stereochemical plausibility and confidence in the model were expressed in the predicted local distance test (plDDT) scores per residue (*49*) and by the Predicted Aligned Error (PAE) measured in Å distance (*50*)(See supplementary Fig 2). For mCXCL4 dimer and tetramer, 5 models were generated with the top-ranking model for each visualised and rendered in PyMol v2.5 with chain-differentiated coloration.

### Biolayer interferometry analysis of CXCL4:GAG binding

An Octet Red96 system (Sartorius AG, Goettingen, Germany) was used with a methodology adapted from Ridley et al. (*11*). GAGs were biotinylated at their reducing end, as previously described (*51*), before immobilisation to High Precision Streptavidin (SAX) biosensors (Sartorius AG, Goettingen, Germany). SAX biosensors were pre-hydrated for 10 mins in assay buffer (10 mM Hepes, 150 mM NaCl, 3 mM EDTA, 0.05% Tween-20, pH 7.4). Immobilisation of heparin dp8 GAG (0.078 µg/ml) in assay buffer was performed to achieve an immobilisation level of approx. 0.05 nm. Sensors were then washed with regeneration buffer (0.1 M Glycine, 1 M NaCl, 0.1% Tween, pH 9.5) before being re-equilibrated in assay buffer. Blank reference or GAG coated sensors were then dipped into chemokines resuspended in 200 µL of assay buffer for 600 sec (association) before being transferred to assay buffer containing wells (dissociation) for at least 600s before surface cleaning with a regeneration buffer wash step. A range of concentrations for human and mouse wild type (50, 25, 12.6, 6.25 and 3.125 nM) and mutant (500, 250, 125, 62.5, 31.25 and 15.625 nM) CXCL4 were analysed. Binding signal was recorded throughout and the signal from binding of chemokine to blank (no immobilised GAG) sensors and by GAG immobilised sensors in assay buffer alone, was subtracted. Data were acquired at 5 Hz and analysed using the Octet HT 10.0 analysis programme to generate association and dissociation constants as well as overall affinity (*K*_D_) value estimates.

### Isolation of mouse brain endothelial cells

Mice were deeply anaesthetised and transcardially perfused with cold PBS. Following perfusion, brains were removed and chopped up before being incubated at 37°C for 30 min with 1 U/ml Liberase TL (Roche) and 40 U/ml DNase I type IV (Sigma) in HBSS H9269 (Sigma). The digestion was stopped with PBS containing 1% FCS (Sigma), with resulting suspension being processed using a manual dounce and passed through a 70 µm cell strainer. Cells were then separated using a 70%/30% Percoll gradient before being washed in PBS.

### Flow Cytometry

Retrieved cells were washed in PBS and stained with Zombie UV fixable cell viability dye (BioLegend, 1:2000 in PBS) for 15 min at room temperature. Cells were next incubated for 10 min at 4°C in 50 µl FcR blocking reagent (BD Biosciences) diluted 1:100. Cells were next washed in flow cytometry buffer (PBS containing 1% fetal bovine serum (FBS, Sigma) and resuspended in 50 µl antibody staining cocktail before being incubated for 30 min at 4°C. Cells were next washed twice in flow cytometry buffer and fixed for 10 min using 1% PFA at room temperature. Cells were then incubated overnight in 50 µl CD31 BV711 (1:100) (Biolegend), CXCL4 Af594 (1:100) (protein foundry) and/or biotinylated anti-heparan sulfate (1:100) (Amsbio). Alternatively, CXCL4 Af594 was pre-incubated with heparin (20 µg) for 60 mins at RT prior to incubation with cells. The next day cells were washed twice with flow cytometry buffer before cells were incubated in 50 µl streptavidin (BV421) for 15 mins at 4°C. Cells were then washed twice in flow cytometry buffer and resuspended in 200 µl flow cytometry buffer before addition of counting beads (ThermoFisher Scientific) and analyzed using a Fortessa flow cytometer (BD Biosciences). Flow cytometry data was analyzed to quantify absolute cell counts or as the percentage of live cells before being normalised relative to vehicle controls to facilitate comparison across experiments.

### Cranial window implantation

Male C57BL/6J mice at around 8 to 12 weeks of age underwent cranial window implantation surgery and intravital imaging was conducted two weeks later, over a period of up to 6 hours. Cranial windows were implanted as previously described (refs below). Animals were placed in an induction chamber with 4% isoflurane and then transferred to a stereotactic frame and maintained at 2-2.5% isoflurane, both in room air. A homeothermic closed loop monitoring system was used to maintain the animal’s body temperature at 37.5 degrees throughout the surgical procedure. The cranium was exposed by removal of a section of skin and the periosteum on top of the head. The wound margin was secured using 3M Vetbond Tissue Adhesive (Thermo Fisher Scientific) and a metal headplate (Narishige CP-2, Japan) was mounted using dental cement (Sun Dental, Japan). A 3 mm biopsy punch was used to gently engrave the outline of the craniotomy onto the skull and the bone was gently thinned around the perimeter using a high-speed micro drill, whilst continuously moving in a circular motion to avoid local heating of the tissue. The surgery area was also regularly soaked with saline to avoid heat damage. The dura was left intact for these experiments.

### Intravital imaging

Anaesthesia was maintained throughout the experiment using 1-2% isoflurane in 100% oxygen. The animal was fixed securely into the imaging setup via the implanted head plate and a homeothermic closed loop monitoring system was used to maintain the animal’s body temperature at 37.5 degrees. Images were collected on a Leica SP8 Upright Multiphoton microscope using a Leica 25x / 0.95 L HC Fluotar dipping objective.

### Intravital imaging of CXCL4 vascular adhesion

Animals underwent intravenous injection of 5 μg CXCL4 594 (ref) and 80 μl of 5 mg/ml FITC dextran (Thermo Fisher Scientific) in sterile saline, following induction of anaesthesia. The MaiTai MP laser (Spectra-Physics) was tuned to 800 nm with emitted signal imaged simultaneously onto external ND-HyD’s through BP525/50 (for Alexa Fluor 488) and BP624/40 (for Texas red) filters. Regions of interest (ROIs) were acquired at a pixel resolution of 1024 x 1024, a frame average of 3 and zoom of 0.75 (equating to a physical size of 620 x 620 μm), with an overall imaging depth of up to 200 μm and a Z step size of 2 μm.

### Lipopolysaccharide treatments

Lipopolysaccharide from *Salmonella enterica* serotype *typhimurium* (LPS, L6511) was purchased from Sigma-Aldrich (St. Louis, MO). LPS was prepared for injection by dissolving in sterile normal saline at a concentration of 0.1mg/ml, and filter sterilizing using a 0.22µm PES filter on the day of use. Mice were weighed and injected intraperitoneally (i.p.) with 0.3mg/kg LPS, or sterile normal saline vehicle once in the morning between the hours of 7:00 and 9:00. *In vivo* transport studies and *in situ* brain perfusions were carried out 27-29 hours post-injection.

### Iodination of mouse CXCL4 and technetium labelling of albumin

Recombinant murine carrier-free wild-type CXCL4 and K56E mutant CXCL4 were purchased from Protein Foundry (Milwaukee, WI). Bovine serum albumin (BSA, 17030-100G) was purchased from Sigma-Aldrich (St. Louis, MO). CXCL4 species were labeled with ^125^I via the chloramine-T method as was described previously (*52*). 20 µg of chemokine was incubated with 1 mCi ^125^I (carrier free, Perkin Elmer) and 10 µg of chloramine-T (Sigma Aldrich) for 1 minute in 0.25 M phosphate buffer. The reaction was terminated by an addition of 100 µg of sodium metabisulfite (Sigma Aldrich), followed by purification on a G-10 Sephadex column. Specific activities were approximately 80Ci/g for both CXCL4 species. Albumin was labelled with ^99m^Tc using the stannous tartrate method as was described previously (*53*). 1 mg albumin was labeled in 500 µl of deionized water with 120-250 µg tin(II) tartrate (S4895) (Sigma-Aldrich, St. Louis, MO) at a pH of 2.5-3.3 with 1 mCi ^99m^Tc. The mixture was incubated for 20 minutes before purification on a Sephadex G-10 column to yield ^99m^Tc-Alb with a specific activity of approximately 100 Ci/g. Protein labelling by iodine and technetium isotopes was characterized by precipitation with 15% trichloroacetic acid (TCA). Greater than 97% radioactivity in the precipitated protein fraction was consistently observed for CXCL4 wild-type and mutant, and greater than 90% radioactivity in the precipitated fraction was consistently observed for ^99m^Tc-Alb. Labelled chemokines were further verified by SDS-PAGE, described below.

### Verification of Iodinated Chemokines by SDS-PAGE

To establish the apparent molecular weight of the unlabelled proteins, 2 µg of CXCL4 wild-type or K56E was run on a pre-cast 4-12% bis-tris PAGE gel in MES buffer (Thermo Fisher, Watham, MA) in reducing or non-reducing conditions as per the manufacturer’s instructions. 5 µl of SeeBlue plus2 protein ladder (LC5925) (Thermo Fisher, Waltham, MA) was used as a size reference. Gels were then stained with SYPRO Ruby Protein Gel Stain (S12001) (Thermo Fisher, Waltham, MA) as per the manufacturer’s instructions. Imaging was performed on an Amersham ImageQuant 800 Biomolecular Imager (General Electeric, Boston, MA) on the UV fluorescence setting with a single exposure.

Following iodination, 100,000 CPM of ^125^I-CXCL4 wild-type and mutant were run on a pre-cast bis-tris PAGE gel as described above for unlabelled chemokines. The gel was then washed twice in deionized water, then incubated in Invitrogen Gel-Dry Drying solution (LC4025) (Thermo Fisher, Waltham, MA) for 15 minutes. The gel was then dried out overnight as per the manufacturer’s instructions. The gel was exposed to autoradiography film (NC9252739) (Thermo Fisher, Waltham, MA) in a light-proof exposure cassette for 24 hours and developed on a Mini-Medical/90 X-ray Film Processor (AFP imaging, Elmsford, NY).

### In vivo measurement of BBB transport

To determine the kinetics of ^125^I-CXCL4 wild-type and mutant brain uptake across the BBB, mice were anesthetized with intraperitoneal urethane and given an intravenous co-injection of 3(10^5^) CPM ^125^I-CXCL4, and 10^6^ CPM ^99m^Tc-Alb (to quantify vascular space) into the left jugular vein. Blood and whole brains were collected after a circulation time from 1 to 30 minutes; in some experiments, mice were perfused with 20 ml lactated Ringer’s solution in about 1 minute to clear blood from the vascular space and to wash away radiotracers that were weakly bound to the vascular lumen. The CPM in brain and serum were measured in a Wizard2 automatic gamma counter (PerkinElmer, Waltham, MA). The amount of radioactivity in the brain and serum was expressed as the percent of injected CPM/g of brain tissue or per µl of serum, respectively (%Inj/g or %Inj/ul). The brain/serum ratios were then calculated by dividing %Inj/g by %Inj/µl to give units of µl/g. Multiple-time regression analysis (*54–56*) was used to determine the rates of chemokine uptake into the brain. This method plots the brain/serum ratios of the compound of interest against a corrected time parameter, exposure time, which is derived from the equation:

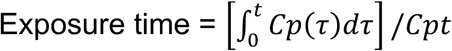

Where t is the time between i.v. injection and blood draw, Cp is the level of radioactivity in serum (expressed as %Inj/ml), Cpt is the level of radioactivity in serum at time t, and *τ* is the dummy variable for time. Exposure time corrects for the clearance of the compound of interest from blood, and so the slope of the linear portion of the brain/serum ratio vs. exposure time relation measures the unidirectional influx rate (Ki, in units of µl/g/min), and the Y-intercept measures Vi (in units of µl/g), which is the vascular space and initial luminal binding at t=0. ^99m^Tc-Alb is a marker of the vascular space, and of any non-specific leakage into tissues that occurs. For ^125^I-CXCL4 uptake calculations, ^99m^Tc-Alb brain/serum ratios were subtracted from the chemokine brain/serum ratios, yielding a ‘delta’ value. The delta values thus correct for unbound chemokines in the residual blood that occupies the vessel lumen and non-specific leakage into the brain.

### In situ measurement of BBB transport

*In situ* brain perfusions were performed as described previously (*57*). Mice were anesthetized with intraperitoneal urethane, and the thorax was opened to expose the heart. Both jugular veins were severed, and the descending thoracic aorta was clamped. A 26-gauge butterfly needle was inserted into the left ventricle of the heart, and Zlokovic’s buffer (7.19 g/l NaCl, 0.3 g/l KCl, 0.28 g/l CaCl_2_, 2.1 g/l NaHCO_3_, 0.16 g/l KH_2_PO_4_, 0.17 g/l anhydrous MgCl_2_, and 0.99 g/l D-glucose, with 10 g/l bovine serum albumin added on the day of perfusion) containing 3(10^4^)/6(10^4^) cpm/ml ^125^I-CCXL4/^99m^Tc-Alb was infused at a rate of 2 ml/min for 1–10 minutes. After perfusion, the mouse was decapitated and the brain was removed and counted in a gamma counter. The perfusion fluid was also counted. Brain/perfusate ratios were calculated by dividing the CPM/g of brain tissue by the CPM/ul of perfusion fluid to yield units of microliters per gram. Because there is no clearance of chemokine or albumin from the perfusate, brain/perfusate ratios were plotted against clock time to determine the Ki and Vi. In some experiments, heparin sodium salt (heparin) (H3393) (Sigma-Aldrich, St. Louis, MO) was added to the perfusate at a concentration of 20 U/ml. In all studies, we verified that the BBB remained intact by co-perfusing ^99m^Tc-Alb. The perfused brain remains viable for approximately 7 hours (*58*), and the BBB remains intact until approximately 12 hours after death (*59, 60*).

### Vascular Depletion in Mice

To determine whether CXCL4 completely crossed the BBB into brain parenchyma and how the percentage of vascular vs. parenchymal CXCL4 changes over time, we performed vascular depletion as adapted to mice (*61, 62*). This method uses a dextran gradient to generate fractions of vascular depleted brain parenchyma and enriched capillaries, which was originally confirmed by measuring the specific activity of y-glutamyl transpeptidase; activity in the parenchymal fraction was 1.91% of that in the pellet, indicating capillary enrichment (*63*). We have consistently found similar enrichment as well by comparing the capillary content of the pellet vs. parenchymal fraction by light microscopy. Mice anesthetized with intraperitoneal urethane received an intravenous injection of 3(10^5^) CPM of ^125^I-CXCL4 (WT or mutant) and 10^6^ CPM of ^99m^Tc-Alb. 30 minutes after intravenous injection, blood from the abdominal aorta was collected. The thorax was then opened, the descending thoracic aorta clamped, the jugular veins severed, and the vascular space of the brain was washed free of blood by perfusing 20 ml of lactated Ringer’s solution through the left ventricle of the heart. The brain was removed, weighed, placed in 0.8 ml of ice-cold physiologic buffer (10 mM HEPES, 141 mM NaCl, 4 mM KCl, 2.8 mM CaCl_2_, 1 mM MgSO_4_, 1 mM NaH_2_PO_4_, and 10 mM D-glucose adjusted to pH 7.4) and rapidly counted in a gamma counter to quantify the total radioactivity in brain. The brain and buffer was then added to a glass Dounce homogenizer plus 1.6 ml of 26% dextran solution. The brain was homogenized with 7 strokes of pestle “A” followed by 7 strokes of pestle “B”. Homogenates were poured into 12×75 plastic polypropylene tubes and centrifuged at 4200g for 20 minutes at 4°C in a Beckman Allegra 21R centrifuge with a swinging bucket rotor (Beckman Coulter, Inc., Fullerton, CA). The pellet containing the brain vasculature and the supernatant containing the brain parenchyma were carefully separated, and the levels of ^125^I and ^99m^Tc were determined in a gamma counter along with serum. The tissue/serum ratios of the capillary and parenchymal fractions were calculated by determining the proportion of total recovered CPM in the respective fraction, and then multiplying the proportion by the tissue/serum ratio calculated for whole brain.

### Acid Precipitation of ^125^I-CXCL4 and ^99m^Tc-albumin from blood and brain tissues

To determine the percentage of ^125^I-CXCL4 (WT or Mutant) that remains intact in blood and brain over time, we performed acid precipitations of serum and brain homogenates. Serum was collected following intravenous injection of 3(10^5^) CPM ^125^I-CXCL4 (WT or mutant) into the left jugular vein. After a circulation time of 1, 2, 5, 10, 15 or 30 minutes, blood was collected from the abdominal aorta, placed on ice for 30 minutes, and then centrifuged at 3500g for 10 minutes. Processing controls were also included where 10^5^ CPM of ^125^I-chemokine (CXCL4 WT or Mut) was added to the blood or brain collection tube, blood or brain was collected from a non-radioactive mouse, and processed along with the test samples. The processing controls are used to estimate the percent degradation that results ex vivo due to tissue processing. 10 µl of serum was then added to 490µl 1% BSA in lactated Ringer’s, and 500 µl of 30% TCA was added and mixed. Samples were centrifuged at 4500g for 10 minutes at 4°C, and then the pellets and the supernatants were counted in a gamma counter. Brains were collected following in situ brain perfusion of 3(10^4^) cpm/ml ^125^I-CXCL4 (WT or mutant) for 10 minutes, and homogenized in 3mls of 1% BSA in lactated Ringer’s solution plus cOmplete Mini protease inhibitor cocktail (11836153001) (Sigma Aldrich, St. Louis, MO). Processing controls were also included where 10^5^ CPM of ^125^I-CXCL4 (WT or mutant) was added to the brain collection tube, non-radioactive brains were collected and processed along with the test samples. The homogenates were centrifuged at 4500g at 4°C for 10 minutes and then 500µl of supernatant was mixed with 500 µl 30% TCA and centrifuged at 4500g at 4°C for 10 minutes. The pellets and supernatants were then counted in a gamma counter.

### Isolation and Culture of Neurospheres from the Hippocampal Region of Mice Brains

Neurospheres were generated from 4-day-old C57BL/6 donor mice. The entire hippocampus was extracted from the brains and digested in 1 mL of DMEM(Gibco 11995-065) containing 1X trypsin (25300054 Life Technologies) at 37°C for 10 minutes. After incubation, mechanical dissociation was performed using a P1000 pipette until no visible tissue fragments remained. Trypsin was diluted by adding four times the volume of DMEM (Gibco 11995-065). The suspension was then centrifuged at 125 × g for 5 minutes, and the supernatant was discarded. The cell pellet was resuspended in 10 mL of basal medium. The basal medium consisted of 75% DMEM high glucose pyruvate (Thermo Fisher, 11995065) and 25% Ham’s F-12 Nutrient Mix with GlutaMax Supplement (Thermo Fisher, 31765035). Additional supplements included 5 µg/mL of Insulin-Transferrin-Selenium (200X, Thermo Fisher, 41400045), 100 µg/mL transferrin (Sigma, T4382), 20 nM progesterone (Sigma, P7556), and 1X penicillin/streptomycin (Thermo Fisher, 15140122). Cell counting was performed using a hemocytometer (KOVA Glasstic Slide 10).

### Neurosphere Culture

Cells were plated in uncoated 6-well plates at a density of approximately 200,000 cells per well in 3 mL of supplemented medium, which consists of basal medium supplemented with 1X B27 (50X, Gibco, 12587-010), 10 ng/mL fibroblast growth factor (FGF, PeproTech, 100-18B), and 10 ng/mL epidermal growth factor (EGF, PeproTech, AF-100-15). Cultures were maintained at 37°C in a humidified incubator with 5% CO₂. The culture medium was refreshed every three days by removing 2 mL of the old medium and carefully adding 2 mL of fresh, pre-warmed supplemented medium. After two weeks, neurospheres were passaged by mechanical dissociation.

### Neurosphere Passage and Dissociation

Primary neurospheres were dissociated after six days in culture and digested in 1 mL of DMEM(Gibco 11995-065) containing 1Xtrypsin (25300054 Life technologies) at 37°C for 5 minutes to generate secondary spheres. Trypsin was diluted by adding four times the volume of DMEM(Gibco 11995-065). The suspension was then centrifuged at 125 × g for 5 minutes, and the supernatant was discarded. The cell pellet was resuspended in 3 mL of supplemented medium, and cell counting was performed using a hemocytometer (KOVA Glasstic Slide 10).

### Treatment and Imaging of Neurospheres

For the CXCL4 and CXCL4 mutant treatment, cells were seeded at densities of 50,000 cells/mL of supplemented medium, with three wells per condition. The following conditions were tested:

- Control (no treatment)
- High concentration of CXCL4 (500 nM, 4.1 µg/mL)
- High concentration of CXCL4 mutant (500 nM, 4.1 µg/mL)
- Low concentration of CXCL4 (65 nM, 0.51 µg/mL)
- Low concentration of CXCL4 mutant (65 nM, 0.51 µg/mL)

Cells were incubated under these conditions for six days. Imaging was performed using light microscopy (CIML Imaging Platform, ZEISS, France).

### Statistics

Prism 8.0 was used for all statistical calculations (GraphPad Software, Inc. San Diego, CA). Linear regression analysis was used to calculate slopes and intercepts, and the standard errors were reported for the slope and y intercepts. Differences in the slopes and intercepts of two lines were evaluated by ANCOVA. Comparison of means group trends was performed using one-way or two-way ANOVA with Dunnett’s or Sidak’s multiple-comparisons testing, respectively.

## Acknowledgments

The authors acknowledge members of the Erickson lab and Dyer lab for critical discussion. We acknowledge members of the biological services, flow cytometry, biomolecular analysis and bioimaging facilities at the University of Manchester. Diagrams created using Biorender.com.

## Funding

Sir Henry Dale fellowship jointly funded by the Wellcome Trust and Royal Society 218570/Z/19/Z (DD)

Wellcome Trust center grant 203128/A/16/Z (DD)

Wellcome Trust Discovery Research platform 226804/Z/22/Z (TAJ, DD)

Wellcome Trust Immunomatrix in Complex Disease (ICD) PhD studentship 218491/Z/19/Z (IM, KOL)

Veterans Affairs Puget Sound Health Care System ACOS Research Fund 2024-1.

## Author contributions

Conceptualization: RR, MAE, DPD.

Methodology: RRW, ALG, SMG, AJLR, SPG, HLB, FCP, IZM, IS, TAJ, JEP, WAB, RR, MAE, DPD.

Investigation: RRW, ALG, SMG, AJLR, SPG, HLB, FCP, IZM, KOL, IS, TAJ, RR, MAE, DPD

Visualization: RRW, ALG, SMG, AJLR, SPG, HLB, KOL, TAJ, RR, MAE, DPD.

Resources: JEP, RR, MAE, DPD. Funding acquisition: JEP, RR, MAE, DPD.

Project administration: JEP, RR, MAE, DPD, Supervision: JEP, RR, MAE, DPD.

Writing – original draft: MAE, DPD

Writing – review & editing: WAB, RR, MAE, DPD

## Data and materials availability

Any data not explicitly evident in the paper will be available on reasonable request.

