## Supplementary information for "Heparan sulfate glycosaminoglycans mediate CXCL4 (PF4) transport across the blood-brain barrier and effects on neurogenesis"

#### Supplementary Figures

##### Mouse CXCL4 WT 8206 Da

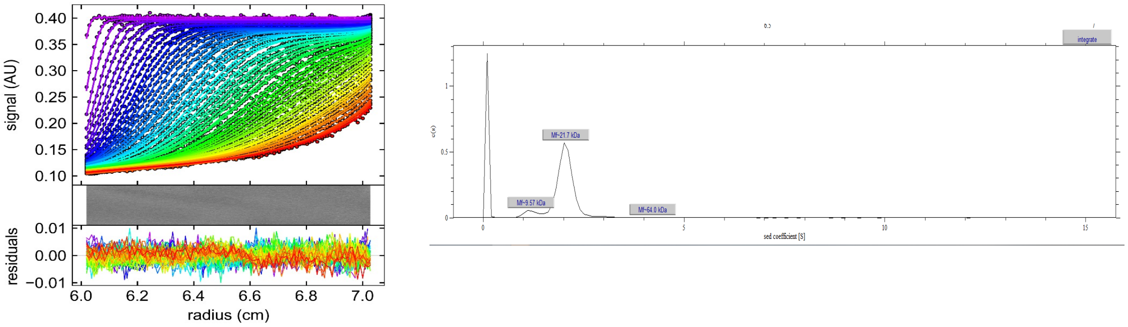


##### Mouse CXCL4 WT 8206 Da + DP8

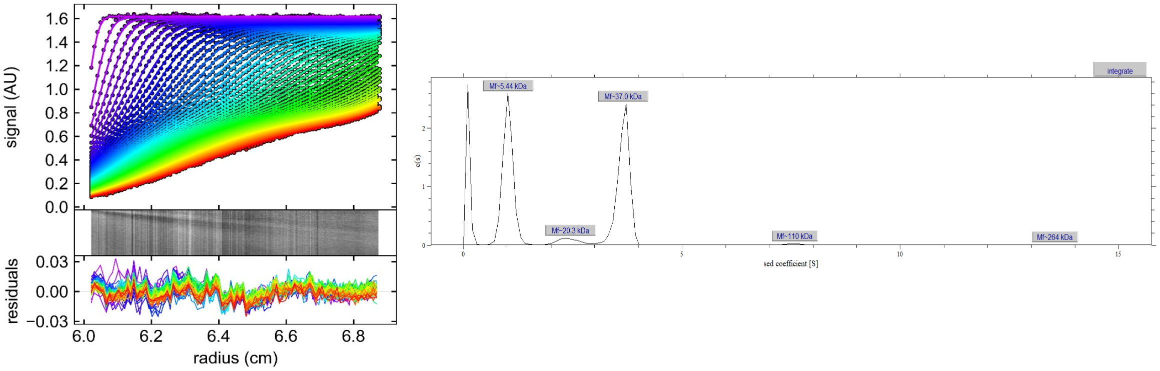


##### Mouse CXCL4 K56E- 8217.7 Da

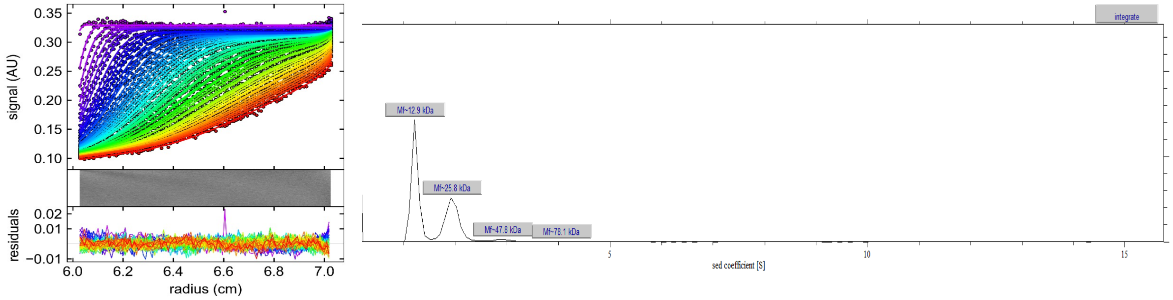


##### Mouse CXCL4 K56E- 8217.7 Da + DP8

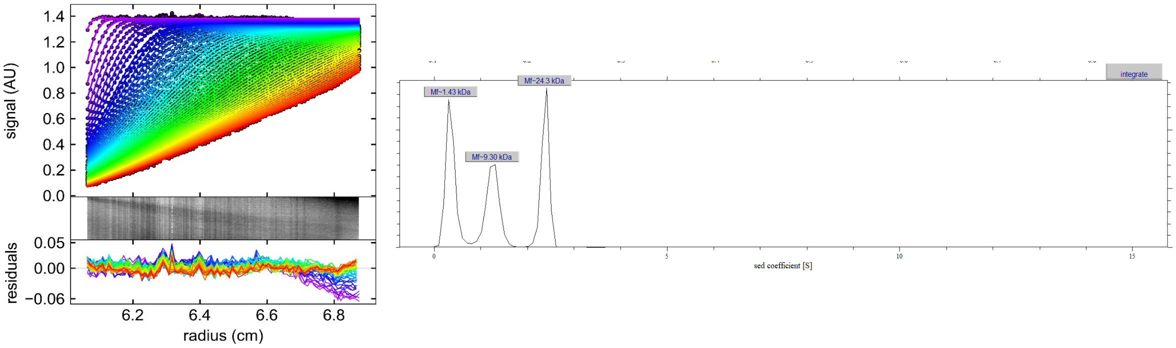

**Supplementary Figure 1. Analytical ultracentrifugation of mouse wild type and mutant CXCL4 with and without GAG.** AUC plots and indicated mass estimates are shown for each condition.

### Mouse CXCL4 WT 8206 Da

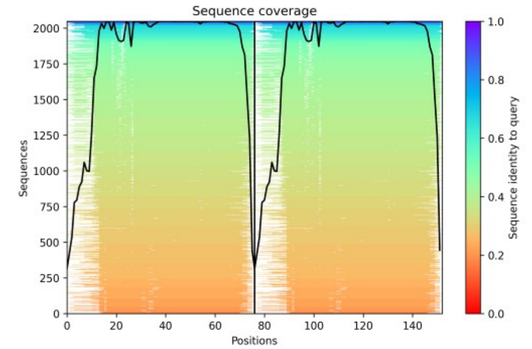


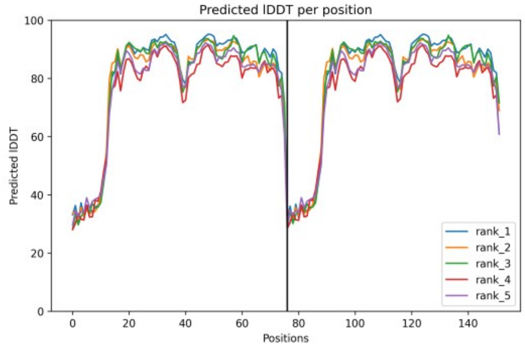


#### PAE scores

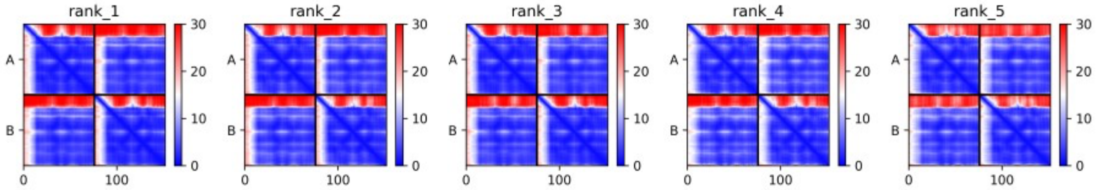


### + dp8 (3.76 s) likely tetramer

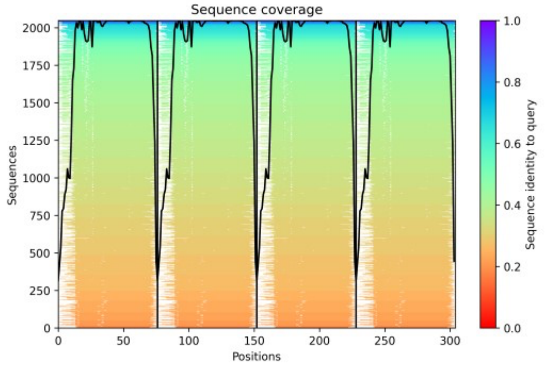


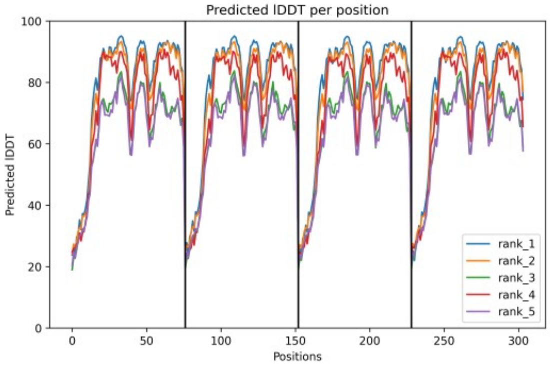


#### PAE scores

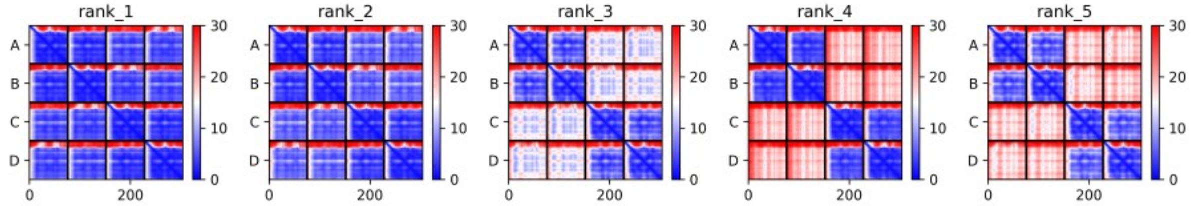

**Supplementary Figure 2: AlphaFold structure prediction of dimeric and tetrameric murine CXCL4.**

Structural predictions of (A-C) dimeric and (D-F) tetrameric mCXCL4 (30-105) were generated by ColabFold: AlphaFold2 multimer model v3. For dimeric mCXCL4, (A) MSA were performed against MMSeqs2 Uniref90 and environmental databases, with the sequence coverage per amino acid position of mCXCL4 displayed. For the 5-most top-ranking models, (B) stereochemical plausibility and confidence in the models were reflected in the plDDT scores per residue position, and (C) PAE scores per residue position represented in Å distance. Tetrameric mCXCL4 was modelled in the same manner as for the dimer, with the corresponding (D) MSA, (E) plDDT and (F) PAE scores for the top 5 ranking models displayed. The top-ranking models for dimeric and tetrameric mCXCL4 were rendered with PyMol v2.5 and are illustrated in Figure 1C.

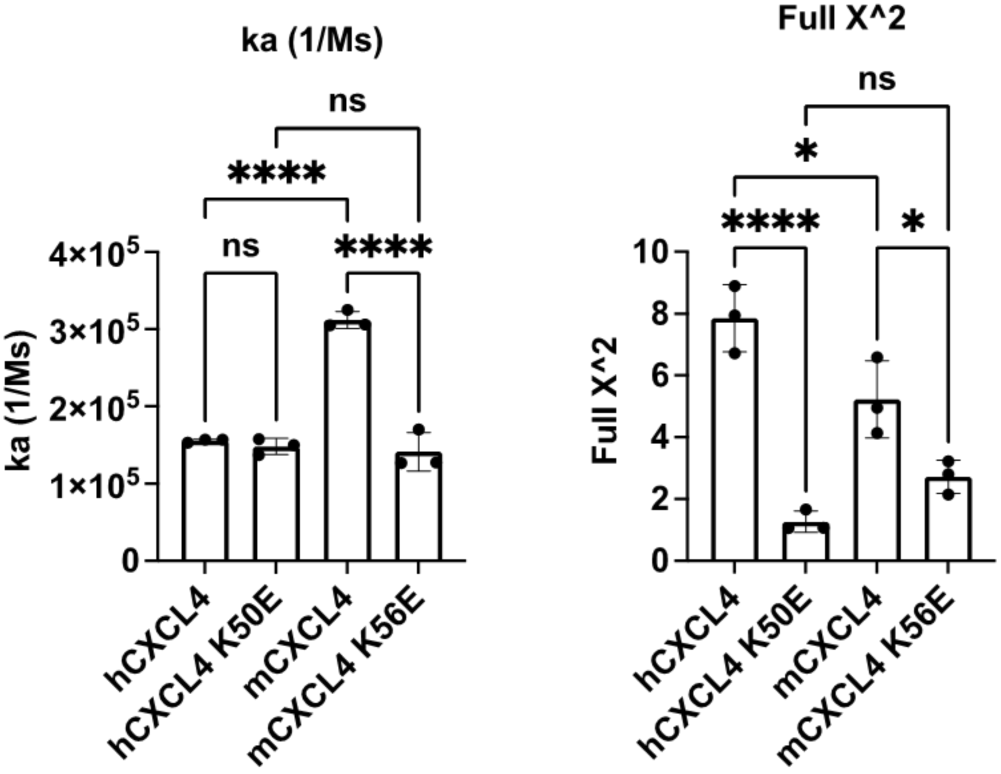


**Supplementary Figure 3. CXCL4:dp8 binding constants.** Association rate constant and curve fitting indication are shown for each condition.

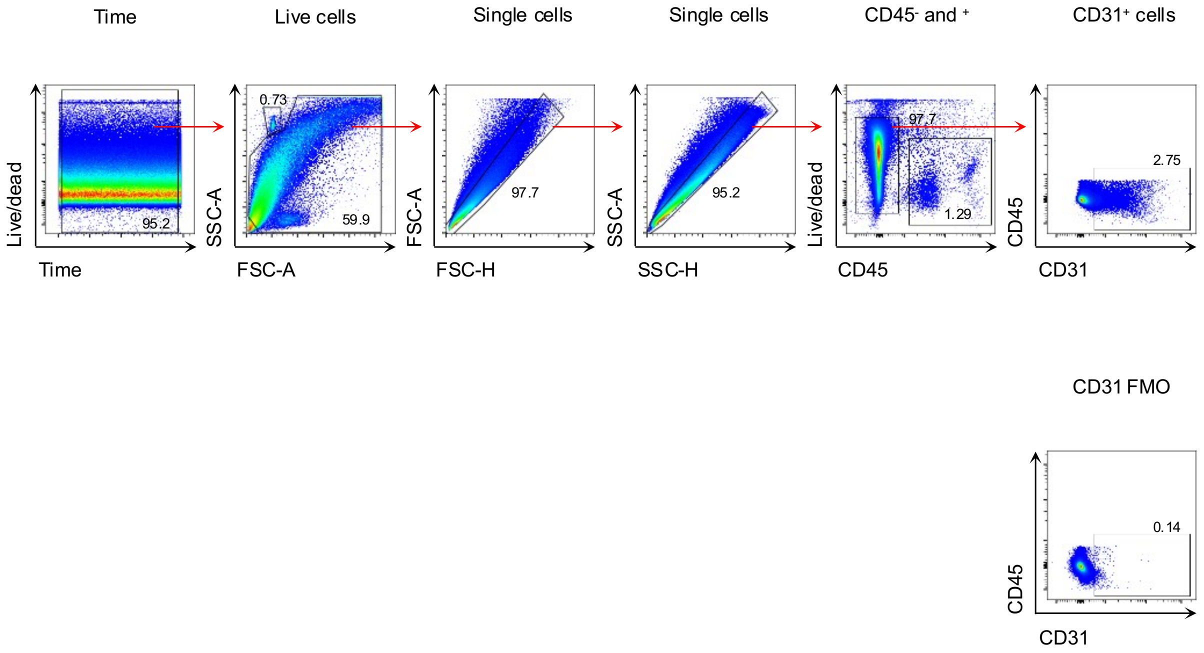


**Supplementary figure 4. Flow cytometry gating for detection of brain endothelial cells (CD45$^{-}$CD31$^{+}$).**

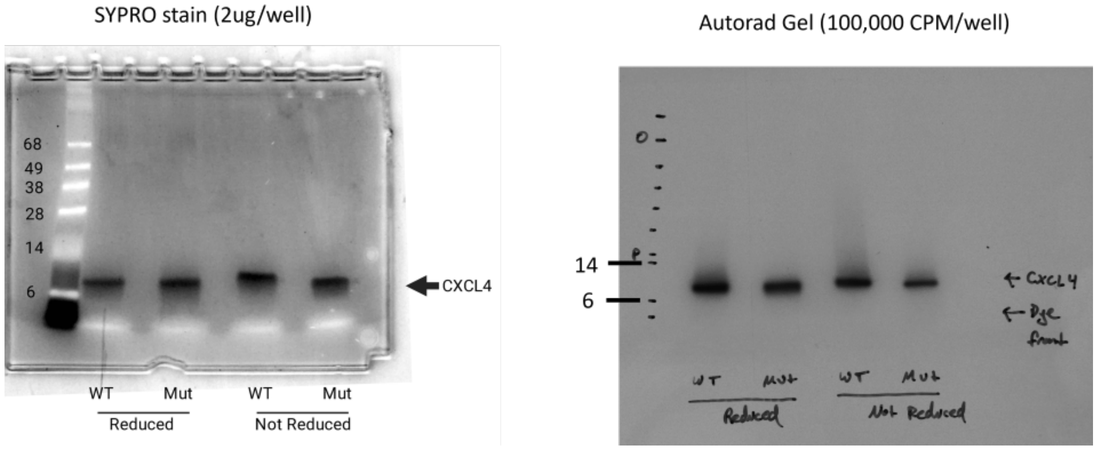


**Supplementary Figure 5: Autoradiography of $^{125}$I-labeled CXCL4** Left panel: Non-radioactive SDS-PAGE of WT and K56E mut CXCL4. Right panel: SDS-PAGE of $^{125}$I-labeled CXCL4 WT and K56E mutant proteins. Proteins were run under reducing and non-reducing conditions.

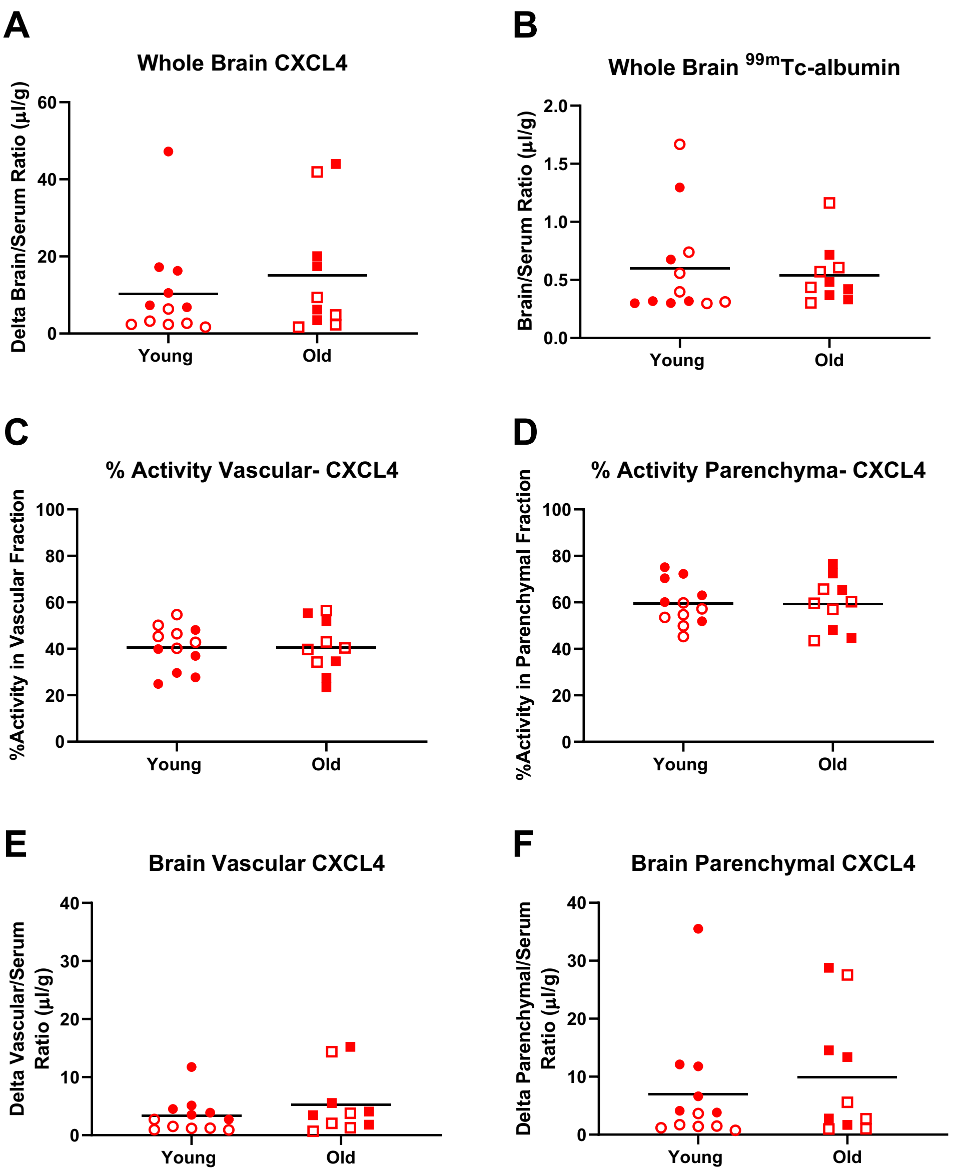


**Supplementary Figure 6: Aging does not affect BBB transport or brain vascular/parenchymal partitioning of $^{125}$I-CXCL4.** Young (2 month) and old (22 month) male (closed shape) and female (open shape) mice (n=10-12/group) were injected i.v. with $^{125}$I-CXCL4 and $^{99m}$Tc-albumin and studied 15 minutes later. Brains were washed out with lactated Ringers and counted, followed by processing for vascular depletion as described in methods. All two-tailed t-tests had $p>0.1$.
